# Sequence-unrelated long noncoding RNAs converged to modulate the activity of conserved epigenetic machineries across kingdoms

**DOI:** 10.1101/2021.02.26.433017

**Authors:** Camille Fonouni-Farde, Aurélie Christ, Thomas Blein, Juan Sebastián Ramírez-Prado, María Florencia Legascue, David Latrasse, Michaël Moison, Leandro Lucero, Lucía Ferrero, Daniel Gonzalez, Moussa Benhamed, Leandro Quadrana, Martin Crespi, Federico Ariel

## Abstract

RNA-DNA hybrid (R-loop)-associated long noncoding RNAs (lncRNAs), including the Arabidopsis lncRNA *AUXIN-REGULATED PROMOTER LOOP* (*APOLO*), are emerging as important regulators of three-dimensional chromatin conformation and gene transcriptional activity. Here, we showed that in addition to the PRC1-component LIKE-HETEROCHROMATIN PROTEIN 1 (LHP1), *APOLO* interacts with the methylcytosine-binding protein VARIANT IN METHYLATION 1 (VIM1), a conserved homolog of the mammalian DNA methylation regulator UBIQUITIN-LIKE CONTAINING PHD AND RING FINGER DOMAINS 1 (UHRF1). The *APOLO*-VIM1-LHP1 complex directly regulates the transcription of the auxin biosynthesis gene *YUCCA2* by dynamically determining DNA methylation and H3K27me3 deposition over its promoter during the plant thermomorphogenic response. Strikingly, we demonstrated that the lncRNA *UHRF1 Protein Associated Transcript* (*UPAT*), a direct interactor of UHRF1 in humans, can be recognized by VIM1 and LHP1 in plant cells, despite the lack of sequence homology between *UPAT* and *APOLO*. In addition, we showed that increased levels of *APOLO* or *UPAT* hamper VIM1 and LHP1 binding to *YUCCA2* promoter. Collectively, our results uncover a new mechanism in which a plant lncRNA coordinates Polycomb action and DNA methylation, and reveal that evolutionary unrelated lncRNAs may exert similar functions across kingdoms.

## INTRODUCTION

In eukaryotes, chromatin structure, composition and dynamics determine the three-dimensional configuration of the genome and are critical for gene regulation, cell fate and function (Yu and Ren, 2017; Doğan and Liu, 2018). Chromatin conformation and related transcriptional states depend on coordinated shifts in DNA methylation, post-translational modifications of histone tails and RNA interference (RNAi) pathways (Tamaru and Selker, 2001; Jaenisch and Bird, 2003; Matzke and Birchler, 2005). In mammalian genomes, DNA methylation is primarily observed at CpG dinucleotides and is estimated to occur at ∼70-80% of CpG sites throughout the genome (Lister et al., 2009). A quarter of non-CG methylation is found in embryonic stem cells, while the remaining unmethylated CpG sites are mostly found in dense clusters, near gene promoters, referred to as CpG islands (Ramsahoye et al., 2000; Suzuki and Bird, 2008; Varley et al., 2013). Unlike in mammals, DNA methylation in plants predominantly occurs on transposons and other repetitive DNA elements, and exists in all possible sequence contexts: symmetric CpG and CpHpG– where H is any base except G – or asymmetric CpHpH (Zhang et al., 2006; Johnson et al., 2007). Non-CpG methylation is mainly distributed at heterochromatin regions, nevertheless, many euchromatic genes exhibit cytosine methylation in their promoter, a feature likely correlated with tissue specificity (Zhang et al., 2006).

In *Arabidopsis thaliana*, the establishment of *de novo* methylation in all sequence contexts is catalyzed by DOMAINS REARRANGED METHYLTRANSFERASE 2 (DRM2), a plant homolog of mammalian DNA (CYTOSINE-5)-METHYLTRANSFERASE 3 (DNMT3) a and b (Cao and Jacobsen, 2002; Law and Jacobsen, 2010). DRM2 is guided to chromatin by 24-nucleotide small interfering RNAs (24nt siRNAs) as part of a pathway known as RNA-directed DNA methylation (RdDM; Cuerda-Gil and Slotkin, 2016). RdDM involves two non-redundant plant specific RNA polymerases, Pol IV and Pol V, in addition to the canonical RNA interference machinery, which requires the activity of RNA-dependent RNA polymerase 2 (RDR2) and members of the Dicer and Argonaute families (Wassenegger et al., 1994; Huettel et al., 2007; Pikaard et al., 2008; Law et al., 2010). Once established, DNA methylation is maintained by three different pathways depending on the sequence context. CpG methylation depends on DNA METHYLTRANSFERASE 1 (MET1), a homolog of mammalian DNA METHYLTRANSFERASE 1 (DNMT1; Finnegan et al., 1996; Kankel et al., 2003). CpHpG methylation relies on CHROMOMETHYLASE 3 (CMT3), a plant specific DNA methyltransferase that associates with SU(VAR)3–9 HOMOLOG (SUVH) histone methyltransferases and recognizes dimethylated histone 3 tails at lysine 9 (H3K9me2; Ebbs and Bender, 2006; Henderson and Jacobsen, 2007; Johnson et al., 2007; Du et al., 2012; Stroud et al., 2014). Finally, CpHpH methylation is maintained through persistent *de novo* methylation by DRM2 and by the CMT3 homolog CHROMOMETHYLASE 2 (CMT2) that specifically reads the H3K9me2 mark (Stroud et al., 2013, 2014; Zemach et al., 2013).

Beside the contribution of methyltransferases, the DNA methylation process involves methylcytosine-binding proteins of the VARIANT IN METHYLATION (VIM/ORTH) family (Woo et al., 2007, 2008). VIM/ORTH proteins are homologous to the mammalian UBIQUITIN-LIKE CONTAINING PHD AND RING FINGER DOMAINS (UHRF) proteins known to regulate cytosine methylation through the recruitment of DNMT1 to target loci (Bostick et al., 2007; Sharif et al., 2007; Kraft et al., 2008; Bronner et al., 2019). In particular, UHRF1 functions as an epigenetic regulator maintaining DNA methylation and histone modifications (Xue et al., 2019), and is stabilized by direct interaction with the long noncoding RNA (lncRNA) *UPAT* (*UHRF1 Protein Associated Transcript*; Taniue et al., 2016). VIM proteins are characterized by the presence of a PHD domain recognizing trimethylated histone 3 tails at lysine 4 (H3K4me3; Li et al., 2006; Peña et al., 2006; Shi et al., 2006; Wysocka et al., 2006), a SRA (SET [Su(var), Enhancer of Zeste, Trithorax] and RING [Really Interesting New Gene] Associated) domain that can associate with methylated DNA (Unoki et al., 2004; Johnson et al., 2007; Woo et al., 2007), and two RING domains conferring ubiquitin E3 ligase activity (Kraft et al., 2008). The Arabidopsis genome encodes five highly similar VIM proteins– named VIM1 to 5 – and a related protein ORTH-LIKE1 (ORL1/VIM6; Kraft et al., 2008). VIM1, VIM2 and VIM3 maintain MET1-mediated cytosine methylation at CpG dinucleotides throughout the genome and have been reported to function entirely redundantly to mediate epigenetic transcriptional silencing in collaboration with MET1 (Woo et al., 2008; Stroud et al., 2013; Shook and Richards, 2014).

Interestingly, aberrant changes in active and repressive histone modifications have been observed in *vim1/2/3* and *met1* mutants, supporting that VIM proteins coordinate DNA methylation and histone modification (Soppe et al., 2002; Tariq et al., 2003; Deleris et al., 2012; Kim et al., 2014). In Arabidopsis, the transcriptionally repressive mark histone H3 lysine 27 trimethylation (H3K27me3) is largely restricted to the transcribed regions of single genes, exhibiting a global anti-correlated distribution with centromeric-enriched DNA methylation (Mathieu et al., 2005; Johnson et al., 2007; Deleris et al., 2012). However, a loss of H3K27me3 was also reported at gene bodies in *met1* mutants and at specific VIM1 targets located in euchromatic regions in the *vim1/2/3* triple mutant (Deleris et al., 2012). Notably, the correlation between DNA hypomethylation and H3K27me3 reduction in the *vim1/2/3* mutation was more prevalent in promoter regions than in transcribed regions (Kim et al., 2014b). Collectively, these observations hint at non-canonical mechanisms linking DNA methylation and repressive histone modifications over specific transcriptionally active loci.

In *A. thaliana*, H3K27me3 is deposited by Polycomb group (PcG) proteins in euchromatic regions containing protein-coding genes and is maintained by the plant Polycomb Repressive Complex 1 (PRC1) component LIKE HETEROCHROMATIN PROTEIN 1 (LHP1), a homolog of mammalian HETEROCHROMATIN PROTEIN 1 (HP1; Gaudin et al., 2001; Hsieh et al., 2003; Turck et al., 2007; Veluchamy et al., 2016). LHP1 capacity to localize and mediate epigenetic repression of PcG target genes was shown to rely on its RNA-binding hinge region, suggesting that LHP1 activity could be modulated by interacting RNAs (Berry et al., 2017). Consistently, it was shown that the lncRNA *APOLO* (*AUXIN-REGULATED PROMOTER LOOP*) can regulate local chromatin conformation by decoying LHP1 away from target loci. *APOLO* directly recognizes multiple distant and independent auxin-related loci across the *A. thaliana* genome by short sequence complementarity and the formation of DNA-RNA duplexes (R-loops; Ariel et al., 2014, 2020).

Here, we demonstrated that in addition to the PRC1 component LHP1, *APOLO* lncRNA interacts *in vivo* with VIM1, the plant homolog of UHRF1, linking Polycomb and DNA methylation machineries. RNA-sequencing analyses of *APOLO* over-expression and *vim1* mutant lines revealed that *APOLO* and VIM1 control a large common set of genes related to the thermomorphogenic response. In particular, the *APOLO*-VIM1-LHP1 complex directly targets the heat-responsive auxin-biosynthetic gene *YUCCA2* and conjointly mediates cytosine methylation and H3K27me3 deposition at its promoter, representing a new epigenetic mechanism regulating the plant response to warm temperatures. Strikingly, we also demonstrate that the lncRNA *UPAT*, known to recognize UHRF1 in humans, can interact with VIM1 and LHP1 in plant cells despite the lack of sequence homology between *UPAT* and *APOLO*. Furthermore, similarly to *APOLO* over-expression, *UPAT* constitutive transcription in Arabidopsis plants precludes LHP1 and VIM1 binding to the *YUCCA2* promoter region. Hence, sequence-unrelated lncRNAs may exert similar molecular functions across kingdoms, integrating the epigenetic regulation of gene expression.

## RESULTS

### Long noncoding RNA *APOLO* associates with the methylcytosine-binding protein VIM1 *in vivo*

In order to investigate the composition of the ribonucleoprotein complexes integrated by the lncRNA *APOLO*, we performed an exploratory Chromatin isolation by RNA purification (ChIRP) followed by protein precipitation and mass spectrometry. We used two independent sets of biotinylated DNA probes against *APOLO* (ODD and EVEN sets) and LacZ-probes as a negative control. Among the proteins identified by at least two positive hits of unique peptides in ODD and EVEN, excluded from LacZ ChIRPs, we found the protein VARIANT IN METHYLATION 1 (VIM1, AT1G57820; (Woo et al., 2007)). Transient expression of *GFP*-*VIM1* in *Nicotiana benthamiana* and *Arabidopsis thaliana* leaves revealed that VIM1 is located in the cell nucleus, in agreement with previous observations (Woo et al., 2007). Moreover, this localization was observed regardless of the co-expression with nuclear-enriched *APOLO* transcripts (**Supplementary Figure 1A-B**). Anti-GFP RNA immunoprecipitation (RIP) in *N. benthamiana* leaves transiently co-transformed with *GFP*-*VIM1* and *APOLO*, or in stably transformed Arabidopsis seedlings over-expressing *GFP-VIM1* (OE *VIM1-1*; **Figure 1A**; *VIM1* levels shown in **Supplementary Figure 2A**), revealed a high enrichment of *APOLO* transcripts in VIM1 immunoprecipitated samples. These results indicate that VIM1 is part of a novel ribonucleoprotein complex integrated by *APOLO* lncRNA.

**Figure 1:**
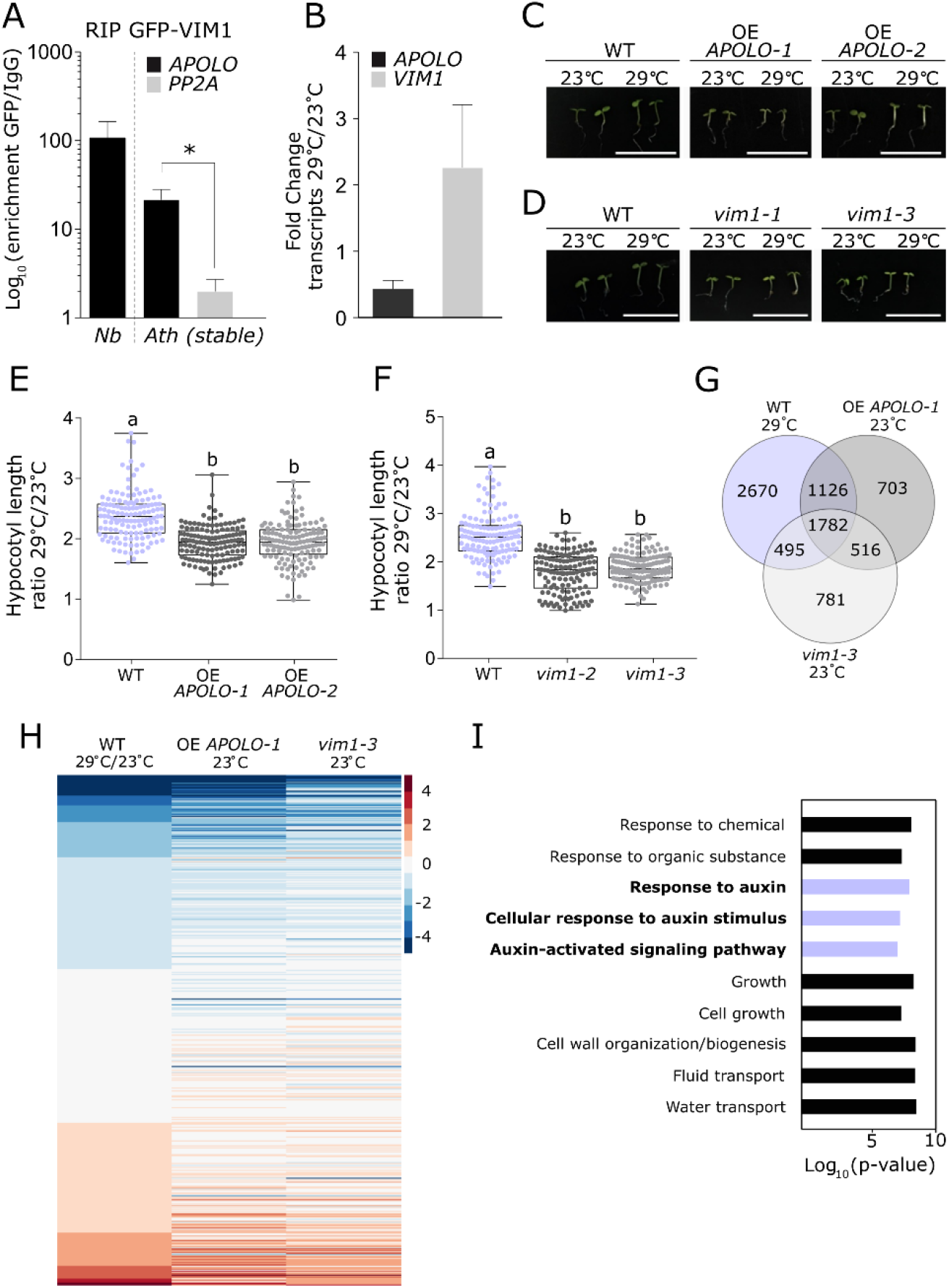
The lncRNA *APOLO* and the methylcytosine-binding protein VIM1 are thermomorphogenesis regulators. (**A**) RNA immunoprecipitation (RIP) assay in *Nicotiana benthamiana* leaves transiently co-transformed with *APOLO* and *GFP-VIM1* translational fusion expressed under the control of the 35S-CaMV promoter, or in *Arabidopsis thaliana* 2-week-old *VIM1* over-expression (OE *VIM1-1*) seedlings. Results are expressed as the log_10_ of the enrichment ratio of the GFP IP over the IgG IP. The non-specific background level of RNA precipitation (*PP2A*) is also shown in Arabidopsis. (**B**) Fold change of *APOLO* and *VIM1* expression levels in relation to 23 °C control conditions in 4-day-old wild-type (WT) seedlings treated with heat (29°C) for 6h. (**C-D**) Representative morphological phenotypes of 4-day-old *APOLO* over-expression (OE *APOLO-1*, OE *APOLO-2*) (in **C**) or *vim1* mutant (*vim1-2*, *vim1-3*) (in **D**) seedlings and their associated WT grown at 23°C or 29°C. Bars, 1cm. (**E-F**) Boxplots showing hypocotyl length quantification ratio at 29°C over 23°C of 4-day-old OE *APOLO-1*, OE *APOLO-2* (**E**) or *vim1-2*, *vim1-3* (**F**) seedlings and their associated WT. Values are represented by colored points. (**G**) Venn diagram of differentially expressed transcripts in WT treated with heat (29°C) for 6h, and in untreated OE *APOLO-1* and *vim1-3* seedlings. (**H**) Heatmap showing log_2_(Fold Change) compared to WT 23°C. A correlation of up- and down-regulated genes in WT in response to 29°C is observed for OE *APOLO-1* and *vim1-3* at 23°C. (**I**) Gene Ontology (GO) enrichment analysis of upregulated transcripts in WT treated with heat. Top ten GO categories with more significant p-values are shown. Violet bars: auxin-related pathways. The ShinyGO Browser is located at bioinformatics.sdstate.edu and published in Ge and Jung (2018). In (**A**), bars represent average ± SD (n = 3 biological replicates). The asterisk indicates significant difference based on a t test (α < 0.05). In (**B**), transcript levels are normalized relatively to the untreated control to show fold changes. Bars represent average ± SD (n = 3 biological replicates). In (**E-F**), results are the mean of three biological replicates and letters indicate significant differences compared to WT, based on a Kruskal-Wallis test (α = 0.05; n ≥ 134).

### *APOLO* and VIM1 are important thermomorphogenesis regulators in *Arabidopsis thaliana*

Abnormal morphological phenotypes, including DNA methylation-dependent late flowering, were previously reported for the *vim1/2/3* triple mutant in contrast to the *vim1* single mutant exhibiting no evident phenotype (Woo et al., 2008). We thus wondered in what developmental context the *APOLO*-VIM1 interaction may occur and exert a regulatory role over target genes. By exploring the eFP Arabidopsis transcriptomic database (Winter et al., 2007), we found that *VIM1*, *VIM2* and *VIM3* genes showed different transcriptional dynamics following heat stress (38°C followed by recovery at 25°C; **Supplementary Figure 3A**, blue box). Therefore, we evaluated the expression of these three genes and *APOLO* at 29°C, a warm temperature usually faced by Arabidopsis in the wild (Parent and Tardieu, 2012). Remarkably, after 6h of exposure to 29°C, *APOLO* transcript levels decreased while *VIM1* transcript levels increased together with the warmth-response markers *PIF4* and *YUC8* (**Figure 1B**; **Supplementary Figure 3B**). In contrast to *VIM1*, the transcriptional accumulation of *VIM2* and *VIM3* was repressed or unaffected, respectively (**Supplementary Figure 3C**).

To assess whether the ribonucleoprotein complex integrated by *APOLO* and *VIM1* is involved in the response to warmth, we tested the effect of their deregulation on the thermomorphogenesis response. Strikingly, two independent *APOLO* overexpression (OE) lines (OE *APOLO-*1 and OE *APOLO-*2; (Ariel et al., 2020)), together with two independent *vim1* insertional mutants resulting in a complete loss of function or knock-down of *VIM1* (*vim1-2* and *vim1-3*, respectively; **Supplementary Figure 2B-C**), exhibited an impaired hypocotyl elongation after four days at 29°C (**Figure 1C-F**; **Supplementary Table 1**). Additionally, we observed a slight but significant reduction of hypocotyl elongation at 29°C only in one out of two independent RNAi *APOLO* lines (RNAi *APOLO-*1 and RNAi *APOLO-*2; Ariel et al., 2014; **Supplementary Figure 4A, C**; **Supplementary Table 1**) and a minor but significant induction of hypocotyl elongation in OE *VIM1* lines (OE *VIM1-*1 and OE *VIM1-*2; **Supplementary Figure 2A**; **Supplementary Figure 4B, D**; **Supplementary Table 1**). Altogether, these results suggest that *APOLO* and VIM1 regulate thermomorphogenesis and incidentally indicate that *VIM1* is not redundant with *VIM2* or *VIM3* in this developmental context.

To pinpoint molecular mechanisms that could explain impaired thermomorphogenesis upon *APOLO* or *VIM1* deregulation, we profiled the transcriptomes of 4-day-old OE *APOLO-*1 and *vim1-3* seedlings grown at 23°C with RNA-Seq of 4-day-old wild-type (WT) seedlings grown at 23°C and subjected or not to heat (29°C) for 6h. In WT seedlings, warmth caused the downregulation of over 3,400 transcripts and the upregulation of approximately 2,650 transcripts (**Supplementary Table 2**). Remarkably, 56% of the total set of up and downregulated transcripts at 29°C in WT were already deregulated at 23°C in OE *APOLO*-1 and/or *vim1-3* (3,403 genes over 6,073; **Figure 1G**), whereas 52% of these transcripts (1,782 out of 3,403) were deregulated in both OE *APOLO*-1 and *vim1-3* at 23°C (**Figure 1G**, central intersection). Moreover, a heatmap analysis revealed that a subset of up and downregulated genes in response to warmth in WT behave consistently in OE *APOLO*-1 and *vim1-3* at ambient temperature (**Figure 1H**). A GO analysis of upregulated transcripts in WT in response to warmth revealed the enriched categories “Response to auxin”, “Cellular response to auxin stimulus” and “Auxin-activated signaling pathway” (**Figure 1H**; **Supplementary Figure 5**), in accordance with the role reported for auxin in thermomorphogenesis (Franklin et al., 2011). Altogether, these transcriptomic analyses hint at a critical role for the *APOLO*-VIM1 complex in the transcriptional reprogramming of gene expression in response to warm temperatures.

We then aimed at identifying the potential direct targets of the *APOLO*-VIM1 complex during the thermomorphogenic response. Given the well-known role of VIM1 in DNA methylation, we compared the list of hypo-methylated genes in *vim1 vs*. WT identified by Bisulfite-Sequencing analyses (BiS-Seq; Stroud et al., 2013), with potential *APOLO* direct targets identified by *APOLO*-ChIRP-Seq (Ariel et al., 2020). Interestingly, among the candidate common targets we found *YUCCA2* (*YUC2*, AT4G13260; **Supplementary Table 3**), a heat-responsive gene involved in auxin biosynthesis (Mashiguchi et al., 2011; Sakata et al., 2010). By using a *proYUC2:GUS* transcriptional fusion (Cheng et al., 2006), we observed that the *YUC2* promoter region is activated in the hypocotyl of 4-day-old seedlings grown at 29°C (**Figure 2A**). In addition, *yuc2* loss-of-function mutant seedlings (Cheng et al., 2006) exhibited a reduced hypocotyl elongation after 4 days at 29°C (**Figure 2B**; **Supplementary Table 1**), indicating that *YUC2* is required for a proper thermomorphogenic response. Consistently, we observed that the transcriptional activation of *YUC2* after 6h at 29°C was impaired in OE *APOLO*-1 and *vim1-3* showing an altered hypocotyl elongation (**Figure 2C**), further suggesting that the *APOLO*-VIM1 complex directly regulates *YUC2* to trigger the plant auxin-related thermomorphogenic response at warm temperatures.

**Figure 2:**
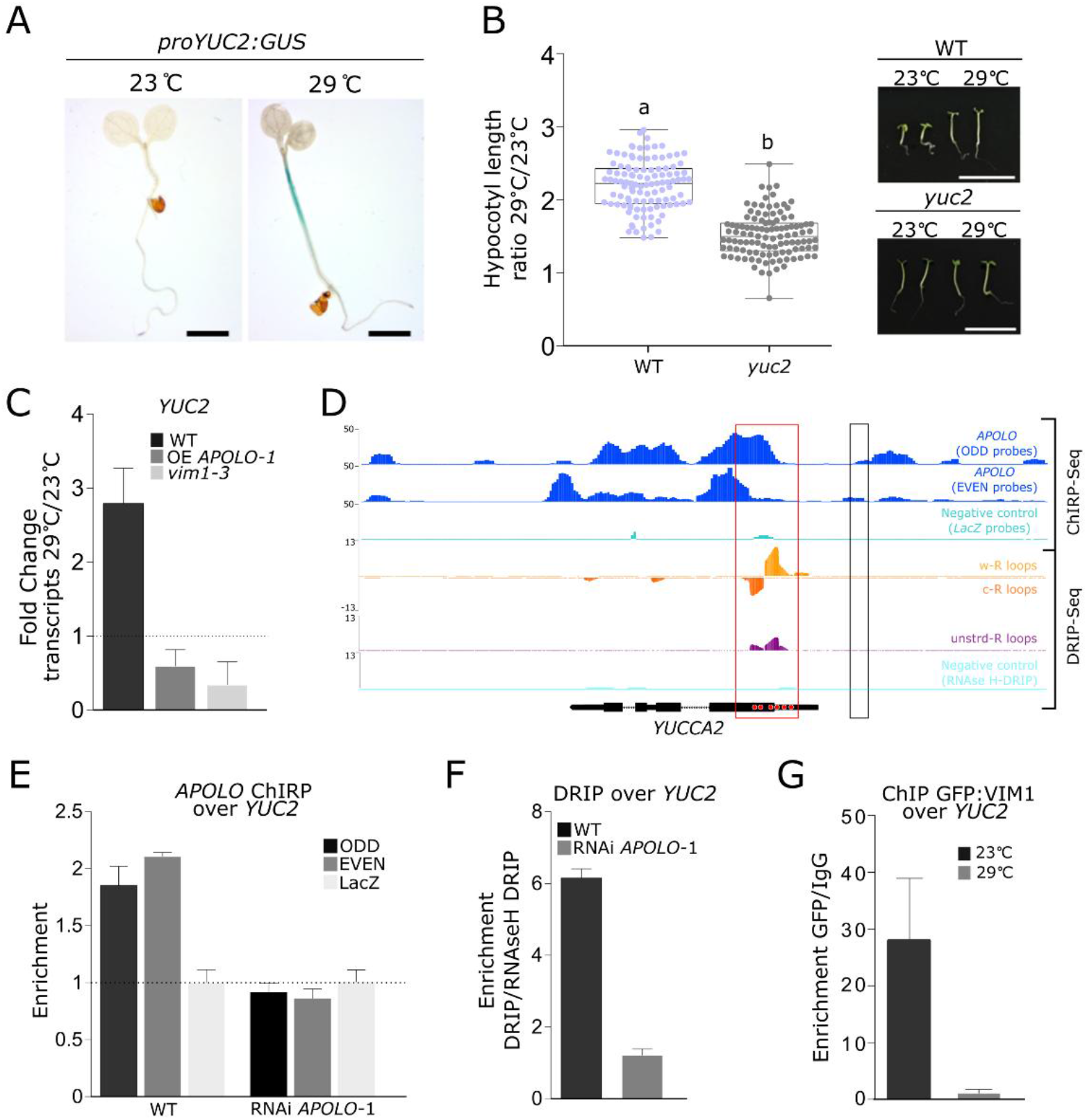
The thermomorphogenesis-related gene *YUCCA2* is directly co-regulated by *APOLO* and VIM1. (**A**) Histochemical localization of GUS activity in 4-day-old seedlings containing the p*YUCCA2:GUS* construct, grown at 23°C or 29°C. Scale bars, 0.1cm. (**B**) Boxplots showing hypocotyl length quantification ratio at 29°C over 23°C of 4-day-old *yuc2* seedlings and their associated wild-type (WT). Values are represented by colored points. Representative morphological phenotypes are shown on the right. Scale bars, 1 cm. (**C**) *YUCCA2* (*YUC2*) transcript levels in 4-days-old WT, OE *APOLO-1* and *vim1-3* seedlings treated or not with heat (29°C) for 6h. (**D**) Epigenetic profile at the *YUC2* locus. Tracks 1 to 3 (Ariel et al. 2020): *APOLO* recognition by chromatin isolation by RNA purification (ChIRP)-Sequencing, using ODD (Track 1) and EVEN (Track 2) sets of probes against *APOLO*. ChIRP negative control using LacZ probes is shown in Track 3. Tracks 4 to 7 (Xu et al. 2017): R-loop formation by DNA:RNA immunoprecipitation (DRIP)-Sequencing, on Watson (Track 4), Crick strand (Track 5) or unstranded sequencing (Track 6). DRIP negative control after RNAseH treatment is shown in Track 7. Gene annotation is shown at the bottom. On the *YUCCA2* schematic representation in the bottom, red dots indicate the presence of six GAAGAA/TTCTTC boxes which may mediate *APOLO* recognition according to Ariel et al. 2020. (**E**) *APOLO* association to DNA of the *YUC2* locus by ChIRP-qPCR in WT and RNAi *APOLO-1* plants. The background level was determined using a set of probes against LacZ RNA. (**F**) RNA-DNA hybrid (R-loop) formation at the *YUC2* locus by DRIP-qPCR in WT and RNAi *APOLO-1* plants. (**G**) Chromatin immunoprecipitation (ChIP)-qPCR analysis of VIM1 binding at the *YUC2* promoter in 4-day-old *VIM1* over-expression (OE *VIM1-1*) seedlings treated or not with heat (29°C) for 6h. In (**A**), one representative picture out of ten stained seedlings is shown. In (**B**), results are the mean of three biological replicates and letters indicate significant differences compared to WT, based on a Mann-Whitney test (α = 0.05; n ≥ 110). In (**C**), transcript levels are normalized relatively to the untreated control to show fold changes. Bars represent average ± SD (n = 3 biological replicates). In (**E-F**), bars represent average ± SD (n = 3 biological replicates). In (**G**), results are expressed as the enrichment ratio of the GFP IP over the IgG IP. Bars represent SD (n = 2 technical replicates). One representative experiment out of two biological replicates is shown.

### *APOLO* and VIM1 directly mediate heat-dependent methylation at the *YUCCA2* promoter

As *APOLO* recognizes its target loci by sequence complementarity and R-loop formation, we first investigated the *APOLO* binding profile over the *YUC2* locus by ChIRP-Seq (Ariel et al., 2020). Compared to the LacZ-probes used as a negative control, ODD and EVEN probes showed specific binding of *APOLO* across the *YUC2* locus, notably to the exon 1, displaying six *APOLO*-binding motifs “GAAGAA/TTCTTC” (**Figure 2D**, first three tracks in blue and light blue, binding motifs indicated as red dots over the *YUC2* locus). *APOLO* ChIRP-qPCR over *YUC2* gene body in WT and RNAi *APOLO-*1 seedlings confirmed the specificity of the interaction (**Figure 2E**). Furthermore, DNA-RNA immunoprecipitation followed by sequencing (DRIP-Seq; Xu et al., 2017) revealed that R-loop formation occurs in the exon 1 of *YUC2*, in correlation with *APOLO* binding (**Figure 2D**, red box), suggesting that *APOLO* directly recognizes *YUC2* through the formation of R-loops. Consistently, DRIP-qPCR in WT and RNAi *APOLO-*1 seedlings revealed that R-loop formation over the *YUC2* locus depends on *APOLO* expression (**Figure 2F**).

Interestingly, a NRPE1 Pol V-subunit Chromatin immunoprecipitation (ChIP)-Seq (Johnson et al., 2014) revealed that the *YUC2* promoter region, located 560 to 1010 bp upstream of *YUC2* TSS, is directly regulated by Pol V in a MET1-dependent manner (**Figure 2D**, black box; **Supplementary Figure 6A**). In addition, a small RNA-Seq profiling reported a temperature-dependent accumulation of RdDM-related 24nt siRNAs over this locus (Gyula et al., 2018; **Supplementary Figure 6B**), suggesting that DNA methylation in this regulatory region may contribute to the temperature-induced transcriptional regulation of *YUC2*. We thus assessed VIM1 binding and DNA methylation at the *YUC2* promoter region in the same developmental stage as our phenotypic characterization. GFP-VIM1 ChIP performed in 4-day-old seedlings grown at 23°C and treated or not at 29°C for 6h, revealed that VIM1 binds to the region located 357 to 494 bp upstream of *YUC2* TSS and that this binding is reduced at 29°C compared to 23°C (**Figure 2G**). In addition, BiS-Seq analyses of WT seedlings grown at 23°C reported cytosine methylation in all sequence contexts in the region located 360 to 610 bp upstream of *YUC2* TSS (**Figure 3A; Supplementary Figure 7**). Remarkably, CpHpH methylation over this region is significantly reduced in *vim1-3* (**Figure 3B**; **Supplementary Figure 7**), hinting at a critical role for VIM1 in the maintenance of non-CpG methylation at the *YUC2* locus. Methylated DNA immunoprecipitation (MeDIP) served to confirm these results and additionally revealed that DNA methylation is reduced in this region in the WT after 6h at 29°C (**Figure 3C**). Furthermore, in OE *APOLO-*1 and *vim1-3*, DNA methylation levels resulted to be lower at 23°C and increased at 29°C, exhibiting the opposite behavior to the WT (**Figure 3C**).

**Figure 3:**
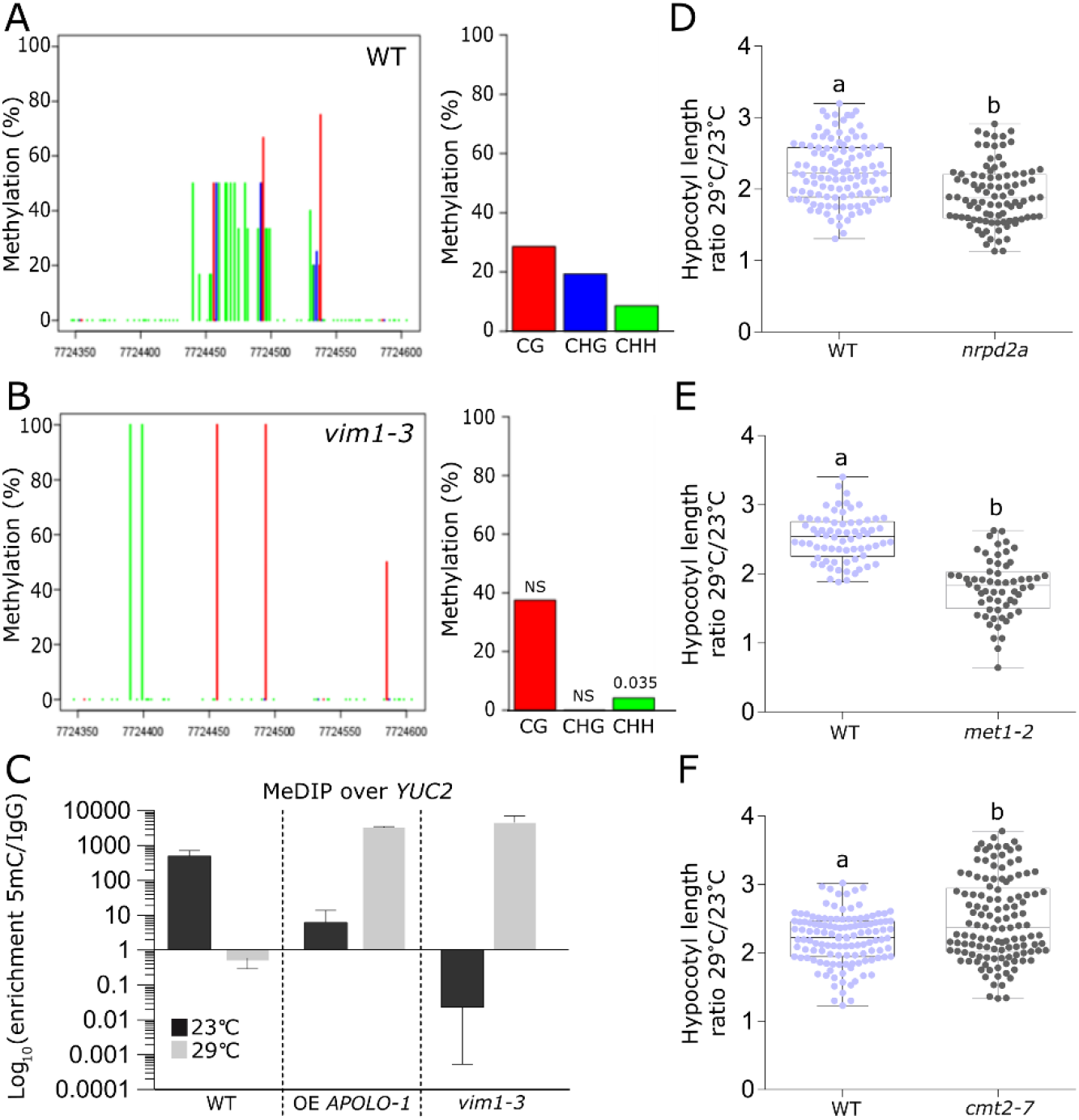
*APOLO* and VIM1 mediate heat-dependent methylation at the *YUCCA2* promoter. (**A-B**) Quantification of DNA methylation at the *YUCCA2* (*YUC2*) promoter (Chr4:7,724,346-7,724,604) by bisulfite (BiS)-Sequencing, in the three sequence contexts CG (red bars), CHG (blue bars) and CHH (green bars), in 4 day-old wild-type (WT) (in **A**) or *vim1-3* (in **B**) mutant seedlings grown at 23°C. (**C**) Methylated DNA immunoprecipitation (MeDIP)-qPCR analysis at the *YUC2* promoter in 4-day-old *APOLO* over-expression (OE *APOLO-1*) or *vim1-3* mutant seedlings treated or not with heat (29°C) for 6h. (**D-F**) Boxplots showing hypocotyl length quantification at 29°C over 23°C of 4-day-old *nrpd2a* (in **D**), *met1-2* (in **E**) or *cmt2-7* (in **F**) mutant seedlings and their associated WT. Values are represented by colored points. In (**A-B**), statistically significant differences between WT and *vim1-3* mutant were calculated using the Fisher’s exact test. NS stands for not significant. In (**C**), results are expressed as the Log_10_ of the enrichment ratio of the 5mC IP over the IgG IP. Bars represent standard deviation (n = 2 biological replicates). In (**D-F**), results are the mean of three biological replicates and letters indicate significant differences compared to WT, based on a Mann-Whitney test (α = 0.05; n ≥ 60).

To further support the relevance of RdDM on *YUC2* regulation in the context of thermomorphogenesis, we additionally characterized the physiological response to warm temperatures of RdDM mutants *nrpd2a* (a common subunit of Pol IV and Pol V), *rdr2-5*, *dcl3-1* and *ago4-8*, as well as CpG methyltransferase mutant *met1*-*2* and non-CpG methyltransferase mutants *cmt2-7* and *cmt3-11* (for a visual summary of their respective roles in RdDM, see **Supplementary Figure 8A**). Hypocotyl elongation at 29°C was reduced in *nrpd2a*, *met1-2, rdr2-5*, *dcl3-1* and *ago4-8* backgrounds, whereas it was unaffected in *cmt3-11* and slightly enhanced in *cmt2-7* backgrounds (**Figure 3D-F**; **Supplementary 8B-H**; **Supplementary Table 1**). In agreement, BiS-Seq analyses (Stroud et al., 2013) revealed that RdDM and *met1-2* mutants showing impaired thermomorphogenesis also displayed reduced CpHpH methylation levels in the *YUC2* regulatory region (**Supplementary Figure 6C-D**, blue box), further supporting that epigenetic modifications regulating DNA methylation at *YUC2* promoter are critical to modulate thermomorphogenesis. Taken together, our results indicate that *APOLO* and VIM1 directly mediate DNA methylation at *YUC2* promoter in response to warm temperatures, in a process likely controlled also by the RdDM pathway.

### VIM1 and LHP1 cooperate to regulate temperature-dependent histone and DNA methylation at the *YUCCA2* promoter

Given that the plant PRC1 component LHP1 recognizes *APOLO in vivo* and co-regulates common target loci across the Arabidopsis genome (Ariel et al., 2014, 2020), we explored whether the *YUC2* locus was regulated by LHP1 and the related transcriptionally repressive mark H3K27me3. ChIP-Seq analyses (Veluchamy et al., 2016) revealed that the *YUC2* locus is enriched in LHP1 and H3K27me3 in standard growing conditions (**Figure 4A**), both drastically reduced at 29°C over the promoter region (**Figure 4A**, black box; **Figure 4B-C**). Moreover, LHP1 binding to the *YUC2* gene body was impaired by *APOLO* over-expression (**Figure 4A**, blue box; **Figure 4D**). Consistent with an involvement of LHP1 in the thermomorphogenic response, 4-day-old *lhp1* loss-of-function mutant seedlings (Veluchamy et al., 2016) exhibited a reduced hypocotyl elongation at 29°C and no induction of *YUC2* after 6h at 29°C (**Figure 4E-F**; **Supplementary Table 1**). Taken together, these results suggest that in addition to VIM1-dependent DNA methylation, the PcG-dependent histone methylation machinery also directly regulates *YUC2* expression in response to warm temperatures.

Considering the similarities between VIM1 and LHP1 behaviors at the *YUC2* locus, we wondered if VIM1-associated DNA methylation and LHP1-related H3K27me3 transcriptional regulations were dependent on each other, and if the two proteins interacted *in vivo*. By performing BiS-Seq of *lhp1 vs.* WT seedlings at 23°C, we observed that the *YUC2* regulatory region in the proximal promoter displayed lower levels of cytosine methylation in all sequence contexts in the *lhp1* mutant (**Figure 5A-B**; **Supplementary Figure 7**, blue box), similarly to the pattern observed in the *vim1-3* mutant (**Figure 3A-B**). Moreover, global DNA methylation resulted to be restored at 29°C in *lhp1*, exhibiting opposite behavior to the WT (**Figure 5C**) and the same trend as OE *APOLO-*1 and *vim1-3* (**Figure 3C**). To further elucidate the link between LHP1 and VIM1, we assessed the levels of H3K27me3 and LHP1 capacity to bind to the *YUC2* promoter region in 4-day-old *vim1-3* seedlings. Notably, LHP1 binding and H3K27me3 levels were impaired in *vim1-3* (**Figure 5D-E**). In combination, these results indicate that while *lhp1* mutation affects DNA methylation of a VIM1-target region, *vim1* mutation impairs LHP1 binding and H3K27me3 deposition on the same locus, hinting at a cooperative interaction between these two epigenetic factors. Additionally, by performing Bimolecular fluorescence complementation (BiFC) assays in *N. benthamiana* leaves transiently transformed with *CYPF*-*VIM1* and *NYFP-LHP1* we observed an interaction between the two proteins (**Figure 5F; Supplementary Figure 9**). Altogether, our results suggest that the lncRNA *APOLO* associates with both a regulator of DNA methylation (VIM1) and a PcG component (LHP1), which further cooperate to mediate DNA methylation and H3K27me3 deposition at a specific *APOLO* target locus.

**Figure 4:**
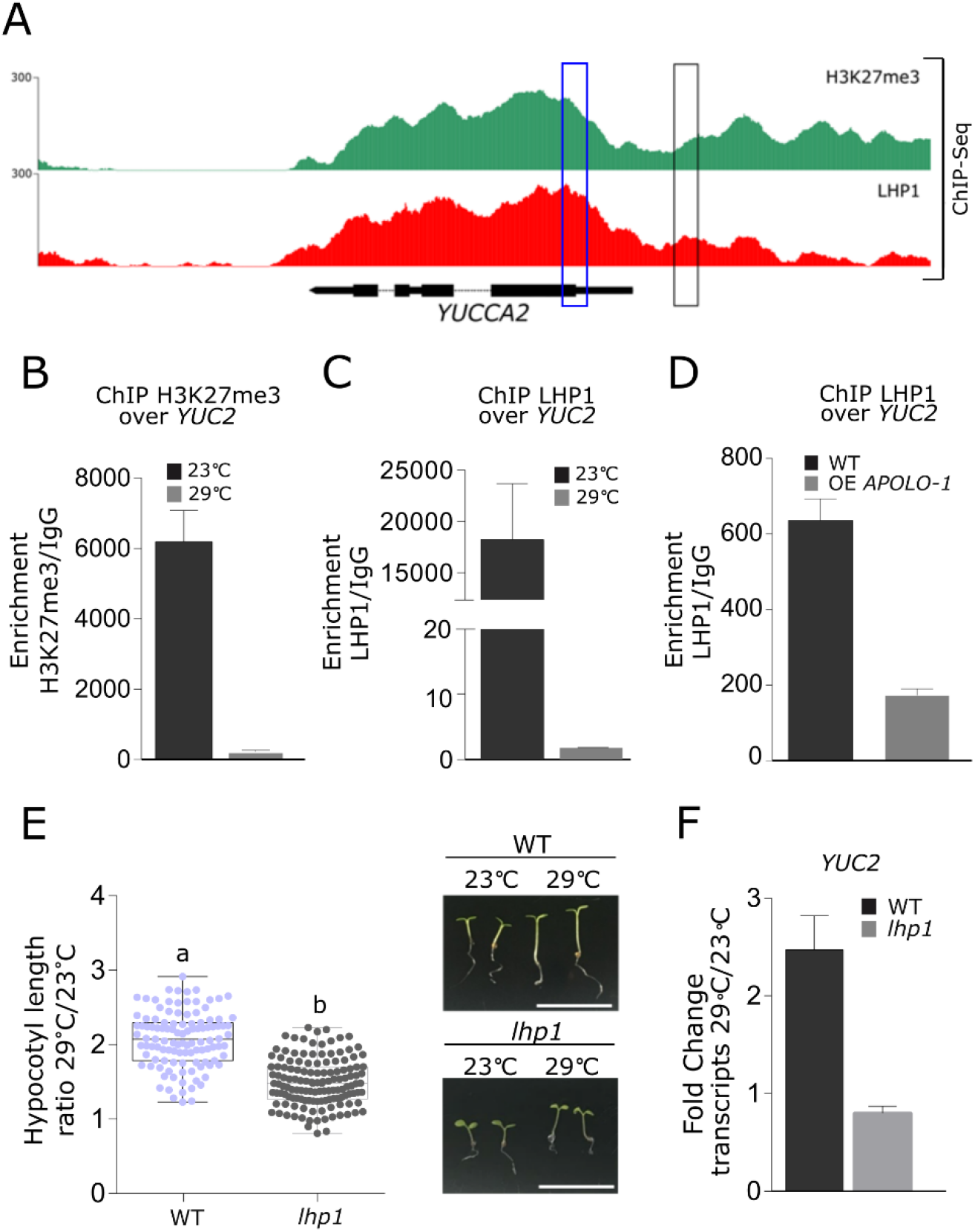
(**A**) Epigenetic landscape at the *YUCCA2* (*YUC2*) locus. Track 1: H3K27me3 deposition by ChIP-sequencing. Track 2: LHP1 binding by ChIP-sequencing (Veluchamy et al. 2016). Gene annotation is shown in the bottom. (**B**) ChIP-qPCR analysis of H3K27me3 deposition at the *YUC2* promoter in 4-day-old wild-type (WT) seedlings treated or not with heat (29°C) for 6h. (**C**) ChIP-qPCR analysis of LHP1 binding at the *YUC2* promoter in 4-day-old WT seedlings treated or not with heat (29°C) for 6h. (**D**) ChIP-qPCR analysis of LHP1 binding at the *YUC2* gene body in WT and *APOLO* over-expression (OE *APOLO-1*) plants. (**E**) Boxplots showing hypocotyl length quantification ratio at 29°C over 23°C of 4-day-old *lhp1* seedlings and their associated WT. Values are represented by colored points. Representative morphological phenotypes are shown on the right. Scale bars, 1 cm. (**F**) *YUC2* transcript levels in 4-days-old WT and *lhp1* seedlings treated or not with heat (29°C) for 6h. In (**B-C**), results are expressed as the enrichment ratio of the H3K27me3 IP (**B**) or LHP1 IP (**C**) over the IgG IP. Bars represent standard deviation (n = 2 technical replicates). One representative experiment out of three biological replicates is shown. In (**D**), bars represent average ± SD (n = 3 biological replicates). In (**E**), results are the mean of three biological replicates and letters indicate significant differences compared to WT, based on a Mann-Whitney test (α = 0.05; n ≥ 111). In (**F**), transcript levels are normalized relatively to the untreated control to show fold changes. Bars represent average ± SD (n = 3 biological replicates).

**Figure 5:**
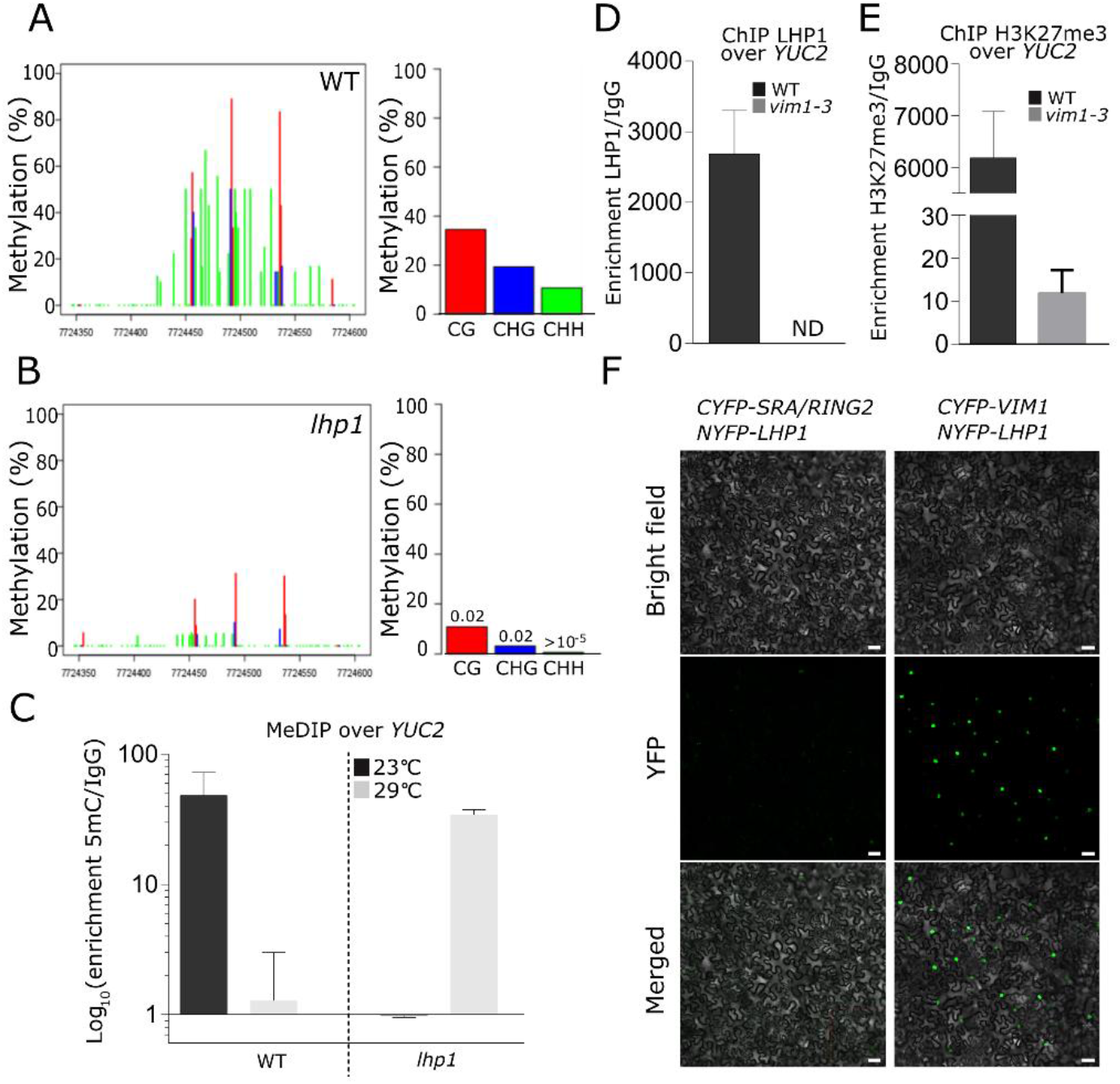
VIM1 and LHP1 co-regulate histone and DNA methylation at the *YUCCA2* promoter. (**A-B**) Quantification of DNA methylation at the *YUCCA2* (*YUC2*) promoter (Chr4:7,724,346-7,724,604) by bisulfite (BiS)-sequencing, in the three sequence contexts CG (red bars), CHG (blue bars) and CHH (green bars), in wild-type (WT) (**A**) or *lhp1* (**B**) mutant seedlings. (**C**) Methylated DNA immunoprecipitation (MeDIP)-qPCR analysis at the *YUC2* promoter in 4-day-old *lhp1* mutant seedlings treated or not with heat (29°C) for 6h. (**D-E**) Chromatin immunoprecipitation (ChIP)-qPCR analyses of LHP1 binding (in **D**) and H3K27me3 deposition (in **E**) at the *YUC2* promoter in 4-day-old WT and *vim1-3* mutant seedlings. ND stands for not detected. (**F**) Bimolecular Fluorescence Complementation (BiFC) assay in transiently transformed *Nicotiana benthamiana* leaves. CYFP was fused to VIM1 or SRA/RING2 and NYFP was fused to LHP1. In both panels, bright-field images (top), YFP fluorescence (middle) and merged images (bottom) are shown. Scale bars, 50μm. In (**A-B**), statistically significant differences between WT and *lhp1* mutant were calculated using the Fisher’s exact test. In (**C**), results are expressed as the enrichment ratio of the 5mC IP over the IgG IP. Bars represent standard deviation (n = 2 biological replicates). In (**D-E**), results are expressed as the enrichment ratio of the LHP1 (in **D**) or H3K27me3 IP (in **E**) over the IgG IP. Bars represent standard deviation (n = 2 technical replicates). One representative experiment out of three biological replicates is shown. In (**F**), one representative picture out of six biological replicates is shown.

### Human lncRNA *UPAT* can exert similar regulatory functions as *APOLO in planta*

Arabidopsis VIM1 is a homolog of the mammalian methylcytosine-binding protein UHRF1 (Kraft et al., 2008; Feng et al., 2010). Interestingly, UHRF1 was shown to directly recognize *in vivo* the human lncRNA *UPAT*, through its flexible spacer region positioned between the SRA and RING domains (Fang et al., 2016; Taniue et al., 2016; Wang et al., 2018). This interaction stabilizes UHRF1 by interfering with its ubiquitination and subsequent degradation (Taniue et al., 2016). In order to determine if VIM1 and UHRF1 interaction with lncRNAs is conserved between plants and animals, we delimited the minimal region of VIM1 that interacts with *APOLO.* We generated independent GFP fusion constructs bearing different *VIM1* coding regions, according to Woo et al., 2007 (**Figure 6A** **right panel**). All the construct-encoded proteins accumulated exclusively or partially in the nucleus of *N. benthamiana* cells, with or without *APOLO* co-expression (**Supplementary Figure 10A-B**). Similar to the reported interaction between UHRF1 and *UPAT* (Taniue et al., 2016), an anti-GFP RIP performed in *N. benthamiana* leaves transiently co-transformed with *APOLO* and GFP-VIM1 derivatives revealed that the two VIM1 portions containing the full spacer region (GFP-SRA/RING2 and GFP-Spacer) bind to *APOLO in vivo*, although with lower efficiency than the full-length VIM1 (**Figure 6A**). Thus, our results indicate that the domain of VIM1 and UHRF1 involved in the interaction with lncRNAs is evolutionary conserved between plants and animals.

**Figure 6:**
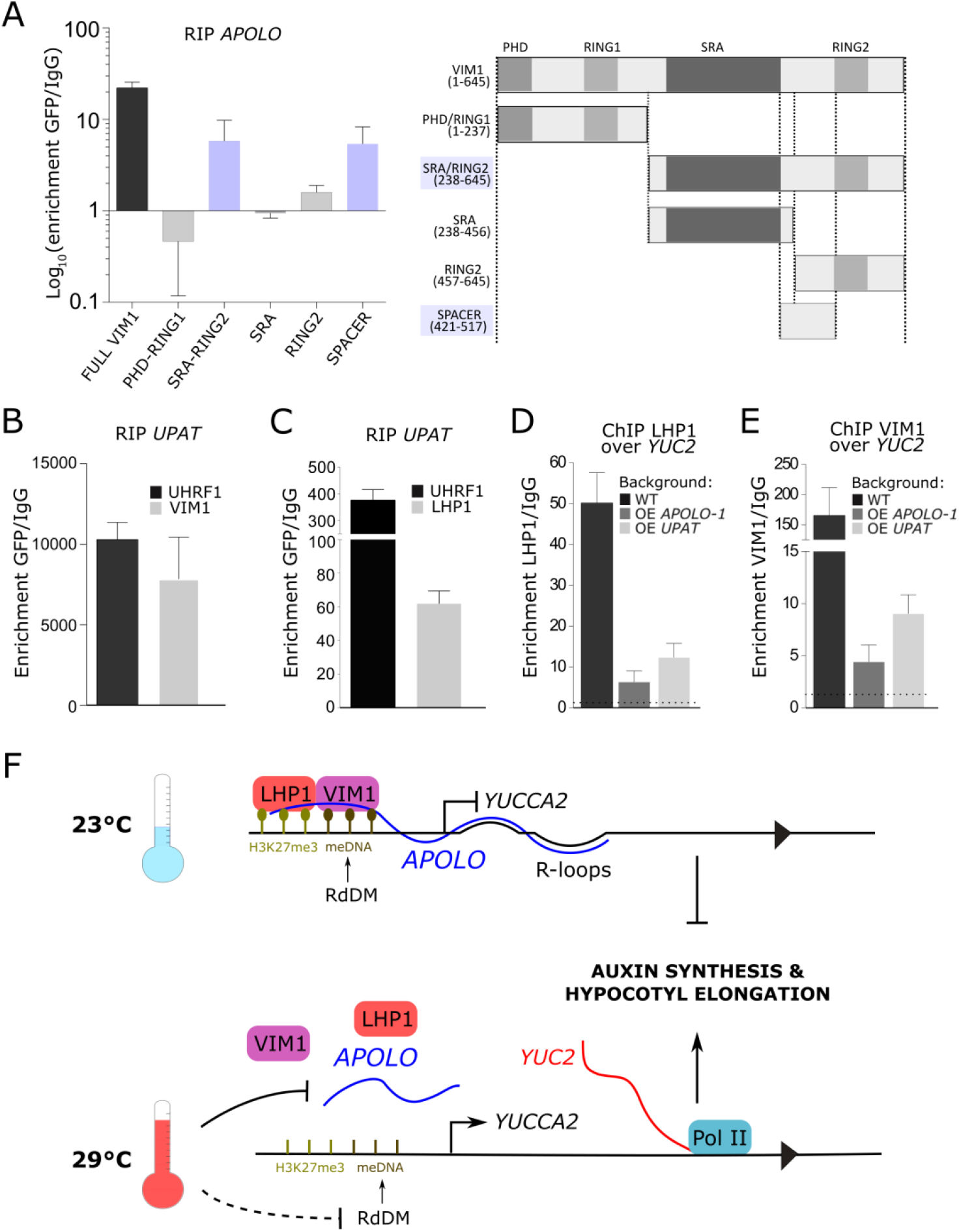
The UHRHF1-interacting lncRNA *UPAT* binds to VIM1 and LHP1 in plant cells and decoys the complex away from chromatin. (**A**) RNA immunoprecipitation (RIP) assay in *Nicotiana benthamiana* leaves transiently co-transformed with *APOLO* and translational fusions expressing *GFP-VIM1* or derivatives (*GFP-PHD*/*RING1*, *GFP-SRA/RING2*, *GFP-SRA*, *GFP-RING2*, *GFP-SPACER*) under the control of the 35S-CaMV promoter. Results are expressed as the Log_10_ of the enrichment ratio of the GFP IP over the IgG IP. Schematic representation of VIM1 protein and derivatives tested for RIP is shown on the right. Amino acid coordinates are indicated between brackets. (**B-C**) RIP assay in *N. benthamiana* leaves transiently co-transformed with *UPAT* and *GFP-UHRF1, GFP-VIM1* (**B**) or *LHP1-GFP* (**C**) translational fusions expressed under the control of the 35S-CaMV promoter. Results are expressed as the enrichment ratio of the GFP IP over the IgG IP. (**D-E**) Chromatin immunoprecipitation (ChIP)-qPCR analysis of LHP1 (in **D**) and VIM1 (in **E**) binding at the *YUC2* promoter in wild-type (WT), *APOLO* over-expression (OE *APOLO-1*) and *UPAT* over-expression (OE *UPAT*) plants transiently transformed or not with *GFP-VIM1*. (**F**) Model for the regulation of *YUC2* expression in response to heat by the *APOLO*-VIM1-LHP1 complex: at 23°C, *APOLO* lncRNA recognizes the *YUCCA2* (*YUC2*) locus by sequence complementarity and R-loop formation. *APOLO* interacts with VIM1 and LHP1 over the *YUC2* promoter region, which exhibits RdDM and H3K27me3 deposition, blocking *YUC2* transcription. meDNA (light brown balls) and H3K27me3 (dark brown balls) are cooperatively maintained by the VIM1-LHP1 complex. At 29°C, *APOLO* transcript levels decrease and VIM1-LHP1 binding to the *YUC2* promoter region is reduced. Conjointly, 24nt siRNAs accumulation is decreased, impairing RdDM over the *YUC2* promoter. As a result, *YUC2* transcriptional activity increases. YUC2 participates in auxin synthesis, promoting hypocotyl elongation in response to heat. In (**A-F**), bars represent average ± SD (n = 3 biological replicates).

Considering the common ability of the conserved VIM1 and UHRF1 proteins to interact with lncRNAs, we wondered whether the lncRNA *UPAT*, exhibiting no apparent sequence homology with *APOLO* (**Supplementary Figure 10C**), could interact with VIM1 and its partner LHP1 *in planta*. Strikingly, an anti-GFP RIP in *N. benthamiana* leaves transiently co-transformed with *UPAT* and *GFP*-*UHRF1*, *GFP*-*VIM1* or *LHP1-GFP* demonstrated that *UPAT* is able to interact with UHRF1, as well as with VIM1 and LHP1 in plant cells (**Figure 6B-C**).

We thus assessed if the constitutive expression of *UPAT* or *APOLO* could modulate VIM1 and LHP1 binding to *YUC2* promoter. ChIP-qPCR analyses performed in WT, *OE APOLO-1* and *OE UPAT* Arabidopsis stable lines transiently transformed or not with *GFP-VIM1* (**Supplementary Figure 1B)**, revealed that both *APOLO* and *UPAT* over-expression precludes the interaction of VIM1 and LHP1 with *YUC2* promoter (**Figure 6D-E**). Furthermore, *UPAT* constitutive expression in stable Arabidopsis seedlings resulted in decreased transcript levels of *YUC2* (**Supplementary Figure 11A-B**). Altogether, our results revealed a functional interaction between sequence-unrelated long noncoding transcripts and key epigenetic regulators, which is likely conserved across kingdoms (**Figure 6F** for a model in plants).

## DISCUSSION

### Epigenetic transcriptional reprogramming in the thermomorphogenic response

When plants are exposed to warmer nonstressful average temperatures, some organs grow and develop at a faster rate without affecting their final dimensions, whereas others suffer morphological or developmental changes. The latest response is known as thermomorphogenesis and includes petiole elongation, leaf hyponasty and auxin-dependent hypocotyl elongation (Gray et al., 1998; Casal and Balasubramanian, 2019; Bellstaedt et al., 2019). These modifications of the plant architecture rely on a deep transcriptional reprogramming determined by dynamic changes of chromatin organization (Kumar and Wigge, 2010; Cortijo et al., 2017; Pajoro et al., 2017; Steffen and Staiger, 2017; Susila et al., 2018; Tasset et al., 2018).

A strong correlation between temperature and DNA methylation was first established in Swedish *A. thaliana* accessions, where higher levels of CpHpH methylation were observed in plants grown at 16°C compared to 10°C (Dubin et al., 2015; Kawakatsu et al., 2016). Interestingly, CpHpH methylation is conjointly regulated by CMT2 and the RdDM pathway (Stroud et al., 2013, 2014; Zemach et al., 2013) and both have been involved in the heat response. It was shown that *cmt2* mutants are more tolerant to heat-stress (Shen et al., 2014) and that the heat-stress response relies on the integrity of the RdDM pathway (Popova et al., 2013; Naydenov et al., 2015). In agreement, we observed that *cmt2* mutants displayed a slight but significantly increased response to warm temperatures in contrast to RdDM mutants exhibiting hypocotyl elongation defects. In addition, we observed that *met1* mutants exhibited the same developmental phenotype as RdDM mutants. Accordingly, it was demonstrated that *met1* knockout causes a loss of DNA methylation, a global loss of Pol V at its normal locations and a Pol V redistribution to sites that become hypermethylated (Johnson et al., 2014). Furthermore, we observed a reduction in CpHpH methylation at the *YUC2* promoter in the *vim1-3* mutant, linking the canonical CpG-related machinery with the RdDM pathway.

Importantly, *met1* was also shown to suffer H3K9 hypermethylation at PcG target genes and a redistribution of H3K27me3 to transposons (Deleris et al., 2012), hinting at a relation between DNA methylation and PRC-dependent histone modifications. Consistently, we uncovered here a new molecular link between the DNA methylation-associated protein VIM1 and the H3K27me3-related protein LHP1, which can interact together and with the lncRNA *APOLO*. An indirect relation between LHP1 and VIM1 had already been described through the histone methyltransferase NtSET1, a common interactor. It was first shown in *Nicotiana tabacum* that NtSET1 directly interacts with LHP1 (Yu et al., 2004) and it was then reported that NtSET1 can bind to AtVIM1 *in planta* (Liu et al., 2007), although no link was established between LHP1 and DNA methylation.

Although VIM1, VIM2 and VIM3 have been reported to function redundantly to mediate epigenetic transcriptional silencing (Woo et al., 2008), we showed here that *vim1* single mutants exhibit particular molecular and associated physiological phenotypes in response to warmth. We related the hypocotyl elongation defect observed in *vim1* to the transcriptional activation of the *YUC2* gene (Sakata et al., 2010; Mashiguchi et al., 2011). *YUC2* regulation in response to heat was first observed in flowers, where heat shock (33°C) repressed its expression and caused plant male sterility (Sakata et al., 2010). At milder temperatures (27°C), *YUC2* transcript levels were reported to increase, in correlation with the demethylation of *YUC2* proximal promoter region targeted by RdDM (Gyula et al., 2018), suggesting that cytosine methylation at the *YUC2* regulatory region is critical for the control of its expression in response to heat. In addition to DNA methylation, a correlation was established between the deposition of H3K27me3 and genes exhibiting either high or low transcription rates under warm temperatures (Sidaway-Lee et al., 2014). Consistently, we observed that the *lhp1* mutant displayed impaired hypocotyl elongation and showed no activation of *YUC2* at 29°C. Interestingly, LHP1 was previously reported to exert a positive role in the regulation of *YUCCA* genes under standard growth conditions. Indeed, *YUC5*, *YUC8* and *YUC9* displayed an abnormal transcriptional activity in the *lhp1* mutant (Rizzardi et al., 2011), correlating with a reduction of H3K27me3, as revealed by genome-wide approaches (Veluchamy et al., 2016). In addition, LHP1 was found to localize to *YUC2*, whose expression was not affected by the *lhp1* mutation at this developmental stage, in an auxin-dependent fashion (Rizzardi et al., 2011).

It was recently shown in Arabidopsis that the 5′ untranslated region of several mRNAs may adopt alternative hairpin conformations under warm cycling daytime temperatures, modulating translation efficiency. It was proposed that mRNA thermoswitches enhancing protein synthesis may constitute a conserved mechanism enabling plants to respond rapidly to high temperatures (Chung et al., 2020). In contrast, the potential role of lncRNAs as thermosensors in higher organisms remains unexplored. In this work, we showed that *APOLO* participates in the thermomorphogenic response in Arabidopsis. Interestingly, *APOLO* transcriptional accumulation increases in response to cold (Moison et al., 2021), whereas we showed here that it decreases under warm temperatures. Further research will be needed to uncover the effect of temperature on the structure of *APOLO* and other lncRNAs that determines their molecular role and contributes to the plant adaptation to environmental constraints.

### R-loop-associated long noncoding RNAs modulate DNA methylation and histone modifications

RNA-DNA hybrids (R-loops) have been identified as important regulators of chromatin conformation and gene transcriptional regulation (Fazzio, 2016; Crossley et al., 2019). *APOLO* is known to recognize its multiple *trans* targets by sequence complementarity and R-loop formation. Upon interaction, it decoys LHP1, shaping local chromatin 3D conformation and subsequently modulating gene transcriptional activity (Ariel et al., 2020). Consistently, we showed here that *APOLO* recognizes *YUC2* gene body through the formation of R-loops.

Interestingly, *APOLO* additionally interacts with VIM1 to co-regulate DNA methylation at the *YUC2* promoter, mediating *YUC2* transcriptional response to warm temperatures. Although the involvement of RNA Pol II-lncRNAs in the regulation of DNA methylation in plants remains largely unknown, previous reports have linked lncRNA activity to promoter methylation in mammals, through the interaction with protein partners (Chalei et al., 2014) or the formation of R-loops (Arab et al., 2014, 2019). Notably, it was reported that DNMT1-interacting lncRNAs can promote or block local DNA methylation (Mohammad et al., 2010; Di Ruscio et al., 2013; Chalei et al., 2014), whereas it was proposed that R-loops formed by nascent transcripts can preclude DNA methylation from promoter regions by repelling DNA methyltransferases (Ginno et al., 2012; Grunseich et al., 2018).

In addition to DNA methylation, a link between R-loops and the activity of PRC complexes has also been proposed. In mammals, it was shown that R-loops can form at a subset of PcG target genes, resulting in alternative regulatory outputs depending on the interplay between the DNA-RNA duplex and PRC components (Skourti-Stathaki et al., 2019). In this work, we showed that the over-accumulation of *APOLO* can titer VIM1 and LHP1 away from the *YUC2* promoter region. It is thus tempting to hypothesize that R-loops formed by *APOLO* at the *YUC2* locus stoichiometrically modulate VIM1 and LHP1 recruitment to DNA in response to changes in ambient temperature. At 23°C, a precise amount of *APOLO* over the *YUC2* locus may guide the LHP1-VIM1 complex to the target region. In contrast, at 29°C *APOLO* transcript levels decrease, hindering VIM1 and LHP1 efficient recognition of *YUC2*. On the other hand, over-accumulation of *APOLO* can decoy LHP1 and VIM1 away from the *YUC2* locus, explaining the similar hypocotyl elongation phenotypes of OE *APOLO*, *vim1* and *lhp1*, as well as the similar DNA methylation levels observed at the *YUC2* promoter in these three backgrounds. The re-methylation of the *YUC2* promoter in OE *APOLO*, *vim1* and *lhp1* at warm temperatures, in contrast to WT, might be due to altered RdDM activity when any component of the regulatory complex is affected. Thereby, we propose that *APOLO* functions as a key mediator of the PRC and DNA methylation machineries, hinting at a stoichiometric factor fine-tuning the activity of R-loops-related lncRNAs.

### Long noncoding RNAs as key mediators of the Polycomb and DNA methylation machineries

Connections between Polycomb and DNA methylation machineries have been reported in few studies in mammals. DNMT1 was shown to directly interact with HP1 to mediate silencing of euchromatic genes (Smallwood et al., 2007) and its enzymatic activity is also stimulated by direct interaction with UHRF1 (Nady et al., 2011; Rothbart et al., 2012; Nishiyama et al., 2013; Bashtrykov et al., 2014; Zhao et al., 2016). Interestingly, HP1 and UHFR1 have been shown to co-exist on chromatin (Rose and Klose, 2014), although their direct interaction has not been proven. Here, we showed that LHP1 and VIM1 interact *in planta*, suggesting that a direct interaction between their homologs HP1 and UHRF1 may occur in mammals. As in mammals, it was shown in Arabidopsis that VIM1 recruits MET1 to hemi-methylated DNA (Feng et al., 2010; Yao et al., 2012; Kim et al., 2014; Shook and Richards, 2014), hinting at the existence of analogous complexes formed by MET1/DNMT1, VIM1/UHRF1 and LHP1/HP1 in plants and animals.

Remarkably, DNMT1, UHRF1 and HP1 were all shown to interact with coding or long noncoding RNAs (Piacentini et al., 2009; Di Ruscio et al., 2013; Taniue et al., 2016), whereas in plants only LHP1 was shown to bind to RNAs *in vitro* (Berry et al., 2017) and *in vivo* (Ariel et al., 2014, 2020). Notably, UHRF1 can interact with the lncRNA *UPAT* whose scaffold function prevents UHRF1 from ubiquitination and proteasomal degradation (Taniue et al., 2016). Knockdown of *UPAT* results in a drastic decrease in the levels of DNMT1 protein and a reduction of hemimethylation at its target genes (Taniue et al., 2016). Here, we showed that *UPAT* can interact with VIM1 and LHP1 in plant cells, and high amounts of *UPAT* can decoy VIM1 and LHP1 from *YUC2* promoter, as the sequence-unrelated transcript *APOLO*.

Although a global anti-correlation is observed in the Arabidopsis genome between DNA methylation and H3K27me3 deposition, characterizing heterochromatic and euchromatic regions respectively (Johnson et al., 2007; Mathieu et al., 2005; Deleris et al., 2012; Antunez-Sanchez et al., 2020), DNA hypomethylation was associated to H3K27me3 reduction at specific genes in *met1* and *vim1/2/3* mutants (Deleris et al., 2012; Kim et al., 2014b). Consistently, our results indicate that *APOLO* lncRNA interacts with LHP1 and VIM1 to regulate *YUC2*, constituting a lncRNA-mediated non-canonical cooperative interaction between Polycomb and DNA methylation machineries. Although other lncRNAs have been linked to thermomorphogenesis or light dependent hypocotyl elongation, their mechanisms of action or target recognition remain unknown (Wang et al., 2014; Severing et al., 2018). Considering that the sequence-unrelated lncRNA *UPAT* mimicked the action of *APOLO* over the chromatin-binding capacity of the VIM1-LHP1 complex, it is likely that additional yet-uncovered lncRNAs exert a similar role as *APOLO* in Arabidopsis.

Altogether, our results hint at a mechanism possibly conserved across kingdoms, which may rely on the structure rather than the sequence of long noncoding transcripts. A deeper knowledge about the molecular basis behind lncRNA-related regulatory mechanisms will likely push back the frontiers of human therapeutics and will allow the design of innovative strategies for sustainable agriculture in a climate change context.

## METHODS

### Data availability

RNA-sequencing data have been deposited in GEO with the accession code XXX. Bisulfite-sequencing data have been deposited in the European Nucleotide Archive (ENA) under project XXX.

### Plant material and growth conditions

All the *Arabidopsis thaliana* lines used were in the Columbia-0 (Col-0) background. *vim1-2* (SALK_050903) and *vim1-3* (SALK_149277c) seeds were obtained from the Arabidopsis Biological Resource Center (http://www.arabidopsis.org/). Mutant plants were genotyped by PCR using specific primers to amplify the endogenous locus and T-DNA borders (primers used are listed in Supplementary Table 4). *VIM1* and *UPAT* over-expression (OE) transgenic plants were generated through *Agrobacterium tumefaciens* (strain EHA105)-mediated transformation (Clough and Bent, 2008). Two independent lines of transformants harboring *GFP-VIM1* or *UPAT* were selected on MS/2 medium supplemented with kanamycin (40µg/mL). *VIM1* expression levels were measured by RT-qPCR (primers used are listed in Supplementary Table 4).

Seeds were surface-sterilized by treatments with 70% EtOH and 5% hypochloryte-1% SDS, washed and stratified at 4°C for 2 days to obtain homogeneous germination. Seedlings were grown at 23°C or 29°C on solid half-strength Murashige and Skoog (MS/2) medium (Duchefa), under long day conditions (16h light, 95µE.m^-2^.sec^-1^/8h dark). For phenotypic characterization of hypocotyl elongation, seedlings were grown for 3h at 23°C and transferred to 29°C or maintained at 23°C for 4 days. Hypocotyl lengths were measured using the ImageJ software. For RNA-sequencing, chromatin immunoprecipitation and methylated DNA immunoprecipitation assays performed in Arabidopsis stable lines, seedlings were grown for 4 days at 23°C and transferred to 29°C or maintained at 23°C for 6h. For chromatin isolation by RNA purification and DNA-RNA Immunoprecipitation, seedlings were grown for 11 days at 23°C.

### Cloning procedures

The entire coding region of *VIM1* (AT1G57820), *VIM1* derivatives (PHD/RING1, SRA/RING2, SRA, RING2; Woo et al., 2007), *LHP1* (AT5G17690) and the lncRNA *UPAT* (Taniue et al., 2016) were amplified by PCR on genomic DNA or pBluescript:UPAT vector (Taniue et al., 2016) respectively, and cloned into the Gateway entry vector pENTR/D-TOPO (Invitrogen). Entry clones were recombined by Gateway technology (LR reaction) into the pK7WGF2, pK7FWG2 or pK7WG2 vectors containing respectively a p35S-GFP-GW, a p35S-GW-GFP or a p35S-GW cassette (http://www.psb.ugent.be/gateway/index.php). pENTR/VIM1, pENTR/SRA/RING2 and pENTR223/LHP1 were recombined by LR reaction into the pGPTVII.Bar-C/NYFP vectors containing respectively a p35S-CYFP or a p35S-NYFP cassette (http://www.psb.ugent.be/gateway/index.php). The spacer of *VIM1* and the entire coding region of *UHFR1* were amplified by PCR on cDNA and pGE-HIS-MBP *UHRF1* plasmid respectively, and cloned into the GreenGate entry vector pGGC000. Destination vectors were constructed associating modules: 35S promoter, GFP, Ubiquitin 10 terminator and pNOS:BastaR,tNOS, according to Lampropoulos et al. (2013). All Gateway constructs were subsequently transformed into the *A. tumefaciens* strain EHA105 and GreenGate constructs into the *A. tumefaciens* strain ASE containing a pSOUP helper plasmid.

### Transient expression and confocal microscopy

Leaves of *Nicotiana benthamiana* were transiently transformed as described in Waadt and Kudla (2008). For RNA immunoprecipitation, *GFP-VIM1* or derivatives (*GFP-PHD/RING1, GFP-SRA/RING2, GFP-SRA, GFP-RING2* and *GFP-Spacer*) were co-transformed with p35S-*APOLO* (Ariel et al., 2020), or *GFP-UHRF1*, *GFP-VIM1* or *LHP1-GFP* were co-transformed with p35S-*UPAT*. For sub-cellular localization, VIM1 protein or derivatives were co-transformed or not with p35S-*APOLO*. For bimolecular fluorescence complementation, *YFPC-VIM1* or *YFPC-SRA/RING2* were co-transformed with *YFPN-LHP1*. Leaves were harvested or cells were analyzed 2 days after infiltration using a Zeiss confocal microscope equipped with Plan-Apochromat 10x/NA 0,45/M27 or Plan-Apochromat 20x/NA 0,8/r M27 US-VIS-IR dry lens. eGFP and eYFP fluorescence was excited with 488 nm and 514 nm argon laser lines and emission recorded between 490nm and 580nm and 520nm and 600nm respectively. Image acquisitions and analyses were performed on the IPS2 Imaging Facility.

Leaves of 3-week-old *Arabidopsis thaliana* were transiently transformed as described in Zhang et al., 2020. Transformation with *GFP-VIM1* was carried out in leaves > 0.5 cm in length, in 10 to 15 plants per genotype (WT, OE *APOLO-1* and OE *UPAT*). Leaves were crosslinked and harvested (for ChiP) or cells were analyzed 3 days after infiltration, using a Leica TCS SP8 confocal laser scanning microscope. eGFP was excited at 488 nm (intensity=8%) and emission recorded between 495nm and 530nm for GFP and 610nm and 670nm (gain 650) for chlorophyll fluorescence. Images were processed using the Fiji software (Schindelin et al., 2012).

### Chromatin isolation by RNA purification and Mass spectrometry

ChIRP-qPCR was performed as previously described (Ariel et al., 2014, 2020). Formaldehyde-crosslinked seedlings were ground and nuclei were isolated and sonicated using a water bath Bioruptor Pico (Diagenode; 30s on/ 30s off pulses, at high intensity for 10 cycles). *APOLO*-associated chromatin was purified using 100pmol of two independent sets of primers (ODD and EVEN) and a negative control (primers matching the LacZ RNA), respectively (Ariel et al., 2014, 2020). Primers used are listed in Supplementary Table 4.

A method adapted from the ChIRP protocol (Chu et al., 2012) was developed to allow identification of plant nuclear proteins bound to specific lncRNAs, as described in Rigo et al., 2020. Briefly, plants were *in vivo* crosslinked and cell nuclei were purified and extracted through sonication. The resulting supernatant was hybridized against biotinylated complementary oligonucleotides that tile the lncRNA of interest and putative lncRNA-containing protein complexes were isolated using magnetic streptavidin beads. Co-purified ribonucleoprotein complexes were eluted and used to purify RNA or proteins, which were later subject to downstream assays for identification and quantification.

#### Crosslinking and ribonucleoprotein complexes purification

For protein extraction, approximately 250 g of 7-day-old Col-0 plants grown on solid half-strength MS medium was irradiated three times with UV using a CROSSLINKERCL-508 (Uvitec) at 0.400 J/ cm^2^. For RNA extraction, 10g of 7-day-old Col-0 plants grown on solid MS/2 medium was crosslinked under vacuum for 15min with 37ml of 1% (v/v) formaldehyde. The reaction was stopped by adding 2.5ml of 2M glycine and seedlings were rinsed with Milli-Q purified water. For both crosslinking methods, 6g of the fixed material was ground in liquid nitrogen (representing 15 ml of plant material ground to fine dust) and added to 50ml tubes with 25 ml of extraction buffer 1– (the nuclei were prepared starting with 30 tubes; buffer 1: 10mM Tris–HCl pH8, 0.4M sucrose, 10mM MgCl_2_, 5mM β-mercaptoethanol, 1ml/30g of sample powder Plant Protease Inhibitor Sigma P9599). The solution was then filtered through Miracloth membrane (Sefar) into a new tube and 5ml of extraction buffer 2 (10mM Tris–HCl pH8, 0.25M sucrose, 10mM MgCl_2_, 5mM β-mercaptoethanol, 1%Triton X-100, 10 µl 100 µl PMSF) was added. The solution was then centrifuged, the supernatant discarded and the pellet was resuspended in 500 µl of extraction buffer 3 (10mM Tris–HCl pH8, 1.7M sucrose, 2mM MgCl_2_, 5mM β-mercaptoethanol, 0.15% Triton X-100, 50µl protease inhibitor) and layered on top of 600 µl of fresh extraction buffer 3 in a new tube. After centrifugation at 13000rpm for 2min at 4°C to pellet the nuclei, the supernatant was discarded and the pellet resuspended in 300µl of nuclei lysis buffer (50mM Tris–HCl pH7, 1% SDS, 10mM EDTA, 1mM DTT, 50µl protease inhibitor, 10µl RNAse inhibitor per tube) to degrade nuclear membranes. Samples were sonicated three times in refrigerated Bioruptor Plus (Diagenode), 30 cycles 30sec ON/30sec OFF in a Diagenode TPX microtube M-50001. After centrifugation, the supernatant was transferred to a new tube and diluted two times volume in hybridization buffer (50mM Tris–HCl pH7, 750 mM NaCl, 1% SDS, 15% formamide, 1mM DTT, 50µl protease inhibitor, 10µl RNAse inhibitor). One hundred pmol of probes against *APOLO* (ODD and EVEN set of probes, Ariel et al., 2014, 2020) and the corresponding negative set against LacZ were added to samples and incubated 4h at 50°C in a thermocycler. Samples were transferred to tubes containing Dynabeads-Streptavidin C1 (Thermo Fisher Scientific) and incubated 1h at 50°C. Then, samples were placed on a magnetic field and washed three times with 1ml of wash buffer (2× SSC, 0.5% SDS, 1 mMDTT, 100µl protease inhibitor). Protein purification samples for protein extraction were DNase-treated according to the manufacturer (Thermo Scientific). After addition of 1.8ml of TCA-acetone (5ml 6.1N TCA + 45ml acetone + 35µl β-mercaptoethanol), samples were incubated overnight at -80°C. After centrifugation at 20000rpm for 20min at 4°C, the supernatant was discarded and 1.8ml of acetone wash buffer (120ml acetone, 84µl β-mercaptoethanol) was added to the samples. Then, samples were incubated 1h at -20°C and centrifuged again at 20000 rpm for 20min at 4°C. The supernatant was discarded, and the dry pellet was used for mass spectrometry analyses.

### DNA-RNA Immunoprecipitation

DNA-RNA Immunoprecipitation (DRIP) was performed as described in Ariel et al., 2020. Non-crosslinked seedlings were used for nuclei purification and samples were sonicated using a water bath Bioruptor Pico (Diagenode; 30s on/ 30s off pulses, at high intensity for 4 cycles). Chromatin samples were incubated with 50 ml of washed Protein G Dynabeads pre-coated with 1mg of S9.6 antibody (Millipore MABE1095) for 16h at 4°C. Samples treated with RNAseH (Invitrogen) for 2h at 37°C were used for DRIP in parallel as a negative control. After washing, DNA was recovered for qPCR over R-loop related loci (primers used are listed in Supplementary Table 4).

### RNA-sequencing analysis

Total RNA was prepared from 4-day-old *A. thaliana* wild-type (WT), OE *APOLO*-1 (Ariel et al., 2014) and *vim1-3* seedlings using the QIAGEN RNeasy plant mini kit and treated with DNase (Fermentas). Three independent biological replicates were produced per genotype. After RNA extraction, libraries were constructed using the Tru-Seq Stranded mRNA Sample Prep kit (Illumina®). Sequencing was carried out at the IPS2. The Illumina NextSeq500 technology was used to perform 75-bp sequencing. A minimum of 15 million of single end reads by sample was generated. RNA-seq preprocessing included trimming library adapters and quality controls were performed with Trimmomatic (Bolger et al., 2014). Paired end reads with Phred Quality Score Qscore > 20 and read length > 30 bases were kept, and ribosomal RNA sequences were removed with SortMeRNA (Kopylova et al., 2012). Processed reads were aligned using STAR with the following arguments: --alignIntronMin 20 --alignIntronMax 3000 --outSAMtype BAM SortedByCoordinate --alignIntronMax 3000 --outSAMtype BAM SortedByCoordinate --alignIntronMax 3000 --outSAMtype BAM SortedByCoordinate. Read overlapping exons per genes were counted using the featureCounts of the subreads package using the GTF annotation files from the Araport11 project (Cheng et al., 2017). Significance of differential gene expression was estimated using DEseq2 (Love et al., 2014) and the FDR correction of the p-value was used during pairwise comparison between genotypes. A gene was declared differentially expressed if its adjusted p-value (FDR) was < 0.01. Heatmap was generated using log2 transforfed fold change values compared WT 23°C and computed with the pheatmap R (Kolde, 2019).

### Bisulfite-sequencing analysis

Genomic DNA (1μg) was prepared from 4-day-old WT and *vim1-3*, or 14-day-old WT and *lhp1* seedlings, respectively. For WT vs. *vim1-3*, bisulfite conversion, BS-seq libraries and sequencing (paired-end 100 nt reads) were performed by BGI Tech Solutions (Hong Kong). For WT vs. *lhp1*, bisulfite conversion was performed with the Premium Whole Genome Bisulfite Sequencing kit (Diagenode) and libraries (paired-end 100 nt reads) were sequenced at the IPS2 platform. Adapter and low-quality sequences were trimmed using Trimming Galore v0.3.3. Mapping was performed on TAIR10 genome annotation using Bismark v0.14.2 (Krueger and Andrews, 2011) and the parameters: --bowtie2, -N 1, -p 3 (alignment); --ignore 5 --ignore_r2 5 -- ignore_3prime_r2 1 (methylation extractor). Only uniquely mapping reads were retained. The methylKit package v0.9.4 (Akalin et al., 2012) was used to calculate differential methylation in 100 bp bp non-overlapping windows (DMRs). Significance of calculated differences was determined using Fisher’s exact test and Benjamin-Hochberg (BH) adjustment of p-values (FDR<0.05) and methylation difference cutoffs of 40% for CG, 20% for CHG and 20% for CHH. Differentially methylated windows within 100bp or 20bp of each other were merged to form larger DMRs. Only 100bp windows with at least six cytosines covered by a minimum of 6 (CG and CHG) and 10 (CHH) reads in all libraries were considered.

### RNA immunoprecipitation assay

RNA immunoprecipitation (RIP) assays were performed on transiently transformed *N. benthamiana* leaves as described in Sorenson and Bailey-Serres (2015), or in 2-week-old *A. thaliana* OE *VIM1.1* seedlings as described in Bardou et al. (2014), using anti GFP (Abcam ab290) and anti-IgG (Abcam ab6702). RIP was performed using Invitrogen Protein A Dynabeads. Precipitated RNAs were prepared using TRI Reagent (Sigma-Aldrich), treated with DNase (Fermentas) and subjected to RT-qPCR (High Capacity cDNA Reverse Transcription Kit (Thermo); primers used are listed in Supplementary Table 4). Total RNAs were processed in parallel and considered as the input sample.

### Chromatin immunoprecipitation assay

Chromatin immunoprecipitation (ChIP) assays were performed using anti GFP (Abcam ab290), anti LHP1 (Covalab pab0923-P), anti H3K27me3 (Diagenode pab-195-050) and anti-IgG (Abcam ab6702), as described in Ariel et al., 2020. Crosslinked chromatin was sonicated using a water bath Bioruptor Pico (Diagenode; 30sec ON/30sec OFF pulses; 10 cycles; high intensity). ChIP was performed using Invitrogen Protein A Dynabeads. Precipitated DNA was recovered using Phenol:Chloroform:Isoamilic Acid (25:24:1; Sigma) and subjected to RT-qPCR (primers used are listed in Supplementary Table 4). Untreated sonicated chromatin was processed in parallel and considered as the input sample.

### Methylated DNA immunopreciptation

Methylated DNA immunoprecipitation (MedIP) assays were performed using anti 5-mC (Diagenode Mab-081-100) and anti-IgG (Abcam ab6702), as described in Nagymihály et al. (2017). Genomic DNA (1µg) was sonicated using a water bath Bioruptor Pico (Diagenode; 30s on/30s off pulses; 4 cycles; high intensity). MedIP was performed using Invitrogen Protein G Dynabeads. Precipitated DNA was recovered using Phenol:Chloroform:Isoamilic Acid (25:24:1; Sigma) and subjected to RT-qPCR (primers used are listed in Supplementary Table 4). Untreated sonicated genomic DNA was processed in parallel and considered as the input sample.

### RT-qPCR

RT-qPCR were performed using the LightCycler 480 SYBR Green I Master Kit on a StepOne Plus apparatus (Applied Biosystems) using standard protocols (40 to 45 cycles, 60°C annealing). *PP2A* (AT1G13320) was used for normalization (primers used are listed in Supplementary Table 4).

### Histochemical staining

Four-day-old *pYUCCA2:GUS* transgenic seedlings (Cheng et al., 2006) grown at 23°C or 29°C were infiltrated with GUS staining solution (0.1M Phosphate buffer pH 7,5; 100mM K_3_Fe(CN)_6_; 100mM K_4_Fe(CN)_6_; 20% Triton X-100; 50mM X-Gluc) under vacuum and subsequently incubated at 37°C for 12h. Stained tissues were cleared in 70% EtOH for 24h at 37°C and observed under a light microscope.

### Statistical analyses

Statistical analyses were performed with non-parametric tests, Mann–Whitney when n=2 independent samples, and Kruskal-Wallis test when n>2 independent samples.

## Acknowledgments

We thank Y. Zaho (University of California) for *yuc2* and *pYUC2:GUS* Arabidopsis seeds; SB. Rothbart (Van Andel Research Institute) for the pGE-HIS-MBP UHRF1 clone; T. Akiyma (University of Tokio) for the *pBluescript:UPAT* clone. This project was financially supported by grants from Agencia Nacional de Promoción Científica y Tecnológica (PICT), Universidad Nacional del Litoral (CAI+D) and Fima Leloir Award (Argentina); International Associated Lab NOCOSYM (CNRS-CONICET); the Centre National de la Recherche Scientifique (MOMENTUM program, to LQ); IPS2 benefits from the support of Saclay Plant Sciences-SPS (ANR-17-EUR-0007). LL, DG and FA are members of CONICET. CFF, MFL, MM, and LF are fellows of the same institution.

## Author contributions

FA and MC conceived the project; FA, CFF, MC, LQ and MB designed the experiments; CFF, AC, JSRP, DL and FA performed the experiments; LQ, TB and LL analyzed the data; MFL, MM, LF, DG and MB contributed materials and methods; CFF and FA wrote the paper.

## Declaration of interests

The authors declare no competing interests.

## Notes

### Competing Interest Statement

The authors have declared no competing interest.

## References

1. Akalin, A., Kormaksson, M., Li, S., Garrett-Bakelman, F.E., Figueroa, M.E., Melnick, A., and Mason, C.E. (2012). MethylKit: a comprehensive R package for the analysis of genome-wide DNA methylation profiles. Genome Biol. 13.

2. Antunez-Sanchez, J., Naish, M., Ramirez-Prado, J.S., Ohno, S., Huang, Y., Dawson, A., Opassathian, K., Manza-Mianza, D., Ariel, F., Raynaud, C., et al. (2020). A new role for histone demethylases in the maintenance of plant genome integrity. Elife 9, 1–32.

3. Arab, K., Park, Y.J., Lindroth, A.M., Schäfer, A., Oakes, C., Weichenhan, D., Lukanova, A., Lundin, E., Risch, A., Meister, M., et al. (2014). Long noncoding RNA TARID directs demethylation and activation of the tumor suppressor TCF21 via GADD45A. Mol. Cell 55, 604–614.

4. Arab, K., Karaulanov, E., Musheev, M., Trnka, P., Schäfer, A., Grummt, I., and Niehrs, C. (2019). GADD45A binds R-loops and recruits TET1 to CpG island promoters. Nat. Genet. 51, 217–223.

5. Ariel, F., Jegu, T., Latrasse, D., Romero-Barrios, N., Christ, A., Benhamed, M., and Crespi, M. (2014). Noncoding transcription by alternative rna polymerases dynamically regulates an auxin-driven chromatin loop. Mol. Cell 55, 383–396.

6. Ariel, F., Lucero, L., Christ, A., Mammarella, M.F., Jegu, T., Veluchamy, A., Mariappan, K., Latrasse, D., Blein, T., Liu, C., et al. (2020). R-Loop Mediated trans Action of the APOLO Long Noncoding RNA. Mol. Cell 77, 1055–1065.e4.

7. Bardou, F., Ariel, F., Simpson, C.G., Romero-Barrios, N., Laporte, P., Balzergue, S., Brown, J.W.S., and Crespi, M. (2014). Long Noncoding RNA Modulates Alternative Splicing Regulators in Arabidopsis. Dev. Cell 30, 166–176.

8. Bashtrykov, P., Jankevicius, G., Jurkowska, R.Z., Ragozin, S., and Jeltsch, A. (2014). The UHRF1 protein stimulates the activity and specificity of the maintenance DNA methyltransferase DNMT1 by an allosteric mechanism. J. Biol. Chem. 289, 4106–4115.

9. Bellstaedt, J., Trenner, J., Lippmann, R., Poeschl, Y., Zhang, X., Friml, J., Quint, M., and Delker, C. (2019). A mobile auxin signal connects temperature sensing in cotyledons with growth responses in hypocotyls. Plant Physiol. 180, 757–766.

10. Berry, S., Rosa, S., Howard, M., Bühler, M., and Dean, C. (2017). Disruption of an RNA-binding hinge region abolishes LHP1-mediated epigenetic repression. Genes Dev. 31, 2115–2120.

11. Bolger, A.M., Lohse, M., and Usadel, B. (2014). Trimmomatic: A flexible trimmer for Illumina sequence data. Bioinformatics 30, 2114–2120.

12. Bostick, M., Jong, K.K., Estève, P.O., Clark, A., Pradhan, S., and Jacobsen, S.E. (2007). UHRF1 plays a role in maintaining DNA methylation in mammalian cells. Science (80-.). 317, 1760–1764.

13. Bronner, C., Alhosin, M., Hamiche, A., and Mousli, M. (2019). Coordinated dialogue between UHRF1 and DNMT1 to ensure faithful inheritance of methylated DNA patterns. Genes (Basel). 10.

14. Cao, X., and Jacobsen, S.E. (2002). Role of the Arabidopsis DRM methyltransferases in de novo DNA methylation and gene silencing. Curr. Biol. 12, 1138–1144.

15. Casal, J.J., and Balasubramanian, S. (2019). Thermomorphogenesis. Annu. Rev. Plant Biol. 70, 321– 346.

16. Chalei, V., Sansom, S.N., Kong, L., Lee, S., Montiel, J.F., Vance, K.W., and Ponting, C.P. (2014). The long non-coding RNA Dali is an epigenetic regulator of neural differentiation. Elife 3, 1–24.

17. Cheng, C.-Y., Krishnakumar, V., Chan, A.P., Thibaud-Nissen, F., Schobel, S., and Town, C.D. (2017). Araport11: a complete reannotation of the *Arabidopsis thaliana* reference genome. Plant J. 89, 789–804.

18. Cheng, Y., Dai, X., and Zhao, Y. (2006). Auxin biosynthesis by the YUCCA flavin monooxygenases controls the formation of floral organs and vascular tissues in Arabidopsis. Genes Dev. 20, 1790– 1799.

19. Chu, C., Quinn, J., and Chang, H.Y. (2012). Chromatin isolation by RNA purification (ChIRP). J. Vis. Exp. 3912.

20. Chung, B.Y.W., Balcerowicz, M., Di Antonio, M., Jaeger, K.E., Geng, F., Franaszek, K., Marriott, P., Brierley, I., Firth, A.E., and Wigge, P.A. (2020). An RNA thermoswitch regulates daytime growth in Arabidopsis. Nat. Plants 6, 522–532.

21. Clough, S.J., and Bent, A.F. (2008). Floral dip: A simplified method for Agrobacterium-mediated transformation of Arabidopsis thaliana. Plant J. 16, 735–743.

22. Cortijo, S., Charoensawan, V., Brestovitsky, A., Buning, R., Ravarani, C., Rhodes, D., van Noort, J., Jaeger, K.E., and Wigge, P.A. (2017). Transcriptional Regulation of the Ambient Temperature Response by H2A.Z Nucleosomes and HSF1 Transcription Factors in Arabidopsis. Mol. Plant 10, 1258–1273.

23. Crossley, M.P., Bocek, M., and Cimprich, K.A. (2019). R-Loops as Cellular Regulators and Genomic Threats. Mol. Cell 73, 398–411.

24. Cuerda-Gil, D., and Slotkin, R.K. (2016). Non-canonical RNA-directed DNA methylation. Nat. Plants 2, 16163.

25. Deleris, A., Stroud, H., Bernatavichute, Y., Johnson, E., Klein, G., Schubert, D., and Jacobsen, S.E. (2012). Loss of the DNA Methyltransferase MET1 Induces H3K9 Hypermethylation at PcG Target Genes and Redistribution of H3K27 Trimethylation to Transposons in Arabidopsis thaliana. PLoS Genet. 8.

26. Doğan, E.S., and Liu, C. (2018). Three-dimensional chromatin packing and positioning of plant genomes. Nat. Plants 4, 521–529.

27. Du, J., Zhong, X., Bernatavichute, Y. V., Stroud, H., Feng, S., Caro, E., Vashisht, A.A., Terragni, J., Chin, H.G., Tu, A., et al. (2012). Dual binding of chromomethylase domains to H3K9me2-containing nucleosomes directs DNA methylation in plants. Cell 151, 167–180.

28. Dubin, M.J., Zhang, P., Meng, D., Remigereau, M.S., Osborne, E.J., Casale, F.P., Drewe, P., Kahles, A., Jean, G., Vilhjálmsson, B., et al. (2015). DNA methylation in Arabidopsis has a genetic basis and shows evidence of local adaptation. Elife 4, e05255.

29. Ebbs, M.L., and Bender, J. (2006). Locus-specific control of DNA methylation by the Arabidopsis SUVH5 histone methyltransferase. Plant Cell 18, 1166–1176.

30. Fang, J., Cheng, J., Wang, J., Zhang, Q., Liu, M., Gong, R., Wang, P., Zhang, X., Feng, Y., Lan, W., et al. (2016). Hemi-methylated DNA opens a closed conformation of UHRF1 to facilitate its histone recognition. Nat. Commun. 7, 1–12.

31. Fazzio, T.G. (2016). Regulation of chromatin structure and cell fate by R-loops. Transcription 7, 121–126.

32. Feng, S., Cokus, S.J., Zhang, X., Chen, P.Y., Bostick, M., Goll, M.G., Hetzel, J., Jain, J., Strauss, S.H., Halpern, M.E., et al. (2010). Conservation and divergence of methylation patterning in plants and animals. Proc. Natl. Acad. Sci. U. S. A. 107, 8689–8694.

33. Finnegan, E.J., Peacock, W.J., and Dennis, E.S. (1996). Reduced DNA methylation in Arabidopsis thaliana results in abnormal plant development. Proc. Natl. Acad. Sci. U. S. A. 93, 8449–8454.

34. Franklin, K.A., Lee, S.H., Patel, D., Kumar, S.V., Spartz, A.K., Gu, C., Ye, S., Yu, P., Breen, G., Cohen, J.D., et al. (2011). PHYTOCHROME-INTERACTING FACTOR 4 (PIF4) regulates auxin biosynthesis at high temperature. Proc. Natl. Acad. Sci. U. S. A. 108, 20231–20235.

35. Gaudin, V., Libault, M., Pouteau, S., Juul, T., Zhao, G., Lefebvre, D., and Grandjean, O. (2001). Mutations in LIKE HETEROCHROMATIN PROTEIN 1 affect flowering time and plant architecture in Arabidopsis. Development 128, 4847–4858.

36. Ginno, P.A., Lott, P.L., Christensen, H.C., Korf, I., and Chédin, F. (2012). R-Loop Formation Is a Distinctive Characteristic of Unmethylated Human CpG Island Promoters. Mol. Cell 45, 814–825

37. Gray, W.M., Östin, A., Sandberg, G., Romano, C.P., and Estelle, M. (1998). High temperature promotes auxin-mediated hypocotyl elongation in Arabidopsis. Proc. Natl. Acad. Sci. U. S. A. 95, 7197–7202.

38. Grunseich, C., Wang, I.X., Watts, J.A., Burdick, J.T., Guber, R.D., Zhu, Z., Bruzel, A., Lanman, T., Chen, K., Schindler, A.B., et al. (2018). Senataxin Mutation Reveals How R-Loops Promote Transcription by Blocking DNA Methylation at Gene Promoters. Mol. Cell 69, 426–437.e7.

39. Gyula, P., Baksa, I., Tóth, T., Mohorianu, I., Dalmay, T., and Szittya, G. (2018). Ambient temperature regulates the expression of a small set of sRNAs influencing plant development through NF-YA2 and YUC2. Plant Cell Environ. 41, 2404–2417.

40. Henderson, I.R., and Jacobsen, S.E. (2007). Epigenetic inheritance in plants. Nature 447, 418–424.

41. Hsieh, T.F., Hakim, O., Ohad, N., and Fischer, R.L. (2003). From flour to flower: How Polycomb group proteins influence multiple aspects of plant development. Trends Plant Sci. 8, 439–445.

42. Huettel, B., Kanno, T., Daxinger, L., Bucher, E., van der Winden, J., Matzke, A.J.M., and Matzke, M. (2007). RNA-directed DNA methylation mediated by DRD1 and Pol IVb: A versatile pathway for transcriptional gene silencing in plants. Biochim. Biophys. Acta - Gene Struct. Expr. 1769, 358–374.

43. Jaenisch, R., and Bird, A. (2003). Epigenetic regulation of gene expression: How the genome integrates intrinsic and environmental signals. Nat. Genet. 33, 245–254.

44. Johnson, L.M., Bostick, M., Zhang, X., Kraft, E., Henderson, I., Callis, J., and Jacobsen, S.E. (2007). The SRA Methyl-Cytosine-Binding Domain Links DNA and Histone Methylation. Curr. Biol. 17, 379– 384.

45. Johnson, L.M., Du, J., Hale, C.J., Bischof, S., Feng, S., Chodavarapu, R.K., Zhong, X., Marson, G., Pellegrini, M., Segal, D.J., et al. (2014). SRA/SET domain-containing proteins link RNA polymerase V occupancy to DNA methylation. Nature 507, 124–128.

46. Kankel, M.W., Ramsey, D.E., Stokes, T.L., Flowers, S.K., Haag, J.R., Jeddeloh, J.A., Riddle, N.C., Verbsky, M.L., and Richards, E.J. (2003). Arabidopsis MET1 cytosine methyltransferase mutants. Genetics 163, 1109–1122.

47. Kawakatsu, T., Huang, S. Shan C., Jupe, F., Sasaki, E., Schmitz, R.J.J., Urich, M.A.A., Castanon, R., Nery, J.R.R., Barragan, C., He, Y., et al. (2016). Epigenomic Diversity in a Global Collection of Arabidopsis thaliana Accessions. Cell 166, 492–505.

48. Kim, J., Kim, J.H., Richards, E.J., Chung, K.M., and Woo, H.R. (2014a). Arabidopsis VIM proteins regulate epigenetic silencing by modulating DNA methylation and histone modification in cooperation with MET1. Mol. Plant 7, 1470–1485.

49. Kim, S., Kim, D., Cho, S.W., Kim, J., and Kim, J.S. (2014b). Highly efficient RNA-guided genome editing in human cells via delivery of purified Cas9 ribonucleoproteins. Genome Res. 24, 1012– 1019.

50. Kim, S., Park, H.J., Cui, X., and Zhi, D. (2020). Collective effects of long-range DNA methylations predict gene expressions and estimate phenotypes in cancer. Sci. Rep. 10, 1–12.

51. Kolde, R. (2019). pheatmap: Pretty Heatmaps version 1.0.12 from CRAN.

52. Kopylova, E., Noé, L., and Touzet, H. (2012). SortMeRNA: fast and accurate filtering of ribosomal RNAs in metatranscriptomic data. Bioinformatics 28, 3211–3217.

53. Kraft, E., Bostick, M., Jacobsen, S.E., and Callis, J. (2008). ORTH/VIM proteins that regulate DNA methylation are functional ubiquitin E3 ligases. Plant J. 56, 704–715.

54. Krueger, F., and Andrews, S.R. (2011). Bismark: A flexible aligner and methylation caller for Bisulfite-Seq applications. Bioinformatics 27, 1571–1572.

55. Kumar, S.V., and Wigge, P.A. (2010). H2A.Z-Containing Nucleosomes Mediate the Thermosensory Response in Arabidopsis. Cell 140, 136–147.

56. Lampropoulos, A., Sutikovic, Z., Wenzl, C., Maegele, I., Lohmann, J.U., and Forner, J. (2013). GreenGate - A Novel, Versatile, and Efficient Cloning System for Plant Transgenesis. PLoS One 8, e83043.

57. Law, J.A., and Jacobsen, S.E. (2010). Establishing, maintaining and modifying DNA methylation patterns in plants and animals. Nat. Rev. Genet. 11, 204–220.

58. Law, J.A., Ausin, I., Johnson, L.M., Vashisht, A.A., Zhu, J.K., Wohlschlegel, J.A., and Jacobsen, S.E. (2010). A Protein Complex Required for Polymerase V Transcripts and RNA-Directed DNA Methylation in plants. Curr. Biol. 20, 951–956.

59. Li, H., Ilin, S., Wang, W., Duncan, E.M., Wysocka, J., Allis, C.D., and Patel, D.J. (2006). Molecular basis for site-specific read-out of histone H3K4me3 by the BPTF PHD finger of NURF. Nature 442, 91–95.

60. Lister, R., Pelizzola, M., Dowen, R., Hawkins, Rd., Hon, G., Nery, J., Lee, L., Ye, Z., Ngo, Q., Edsall, L., et al. (2009). Human DNA methylomes at base resolution show widespread epigenomic differences. Nature 462, 315–322.

61. Liu, S., Yu, Y., Ruan, Y., Meyer, D., Wolff, M., Xu, L., Wang, N., Steinmetz, A., and Shen, W.H. (2007). Plant SET- and RING-associated domain proteins in heterochromatinization. Plant J. 52, 914–926.

62. Love, M.I., Huber, W., and Anders, S. (2014). Moderated estimation of fold change and dispersion for RNA-seq data with DESeq2. Genome Biol. 15, 550.

63. Mandin, P., and Johansson, J. (2020). Feeling the heat at the millennium: Thermosensors playing with fire. Mol. Microbiol. 113, 588–592.

64. Mashiguchi, K., Tanaka, K., Sakai, T., Sugawara, S., Kawaide, H., Natsume, M., Hanada, A., Yaeno, T., Shirasu, K., Yao, H., et al. (2011). The main auxin biosynthesis pathway in Arabidopsis. Proc. Natl. Acad. Sci. U. S. A. 108, 18512–18517.

65. Mathieu, O., Probst, A. V., and Paszkowski, J. (2005). Distinct regulation of histone H3 methylation at lysines 27 and 9 by CpG methylation in Arabidopsis. EMBO J. 24, 2783–2791.

66. Matzke, M.A., and Birchler, J.A. (2005). RNAi-mediated pathways in the nucleus. Nat. Rev. Genet. 6, 24–35.

67. Mohammad, F., Mondal, T., Guseva, N., Pandey, G.K., and Kanduri, C. (2010). Kcnq1ot1 noncoding RNA mediates transcriptional gene silencing by interacting with Dnmt1. Development 137, 2493– 2499.

68. Moison, M., Pacheco, J.M., Lucero, L., Fonouni-Farde, C., Rodríguez-Melo, J., Christ, A., Bazin, J., Benhamed, M., Ibañez, F., Crespi, M., et al. (2021). The lncRNA APOLO interacts with the transcription factor WRKY42 to trigger root hair cell expansion in response to cold. BioRxiv 2020.07.13.188763.Molecular Plant (accepted, in press)

69. Nady, N., Lemak, A., Walker, J.R., Avvakumov, G. V., Kareta, M.S., Achour, M., Xue, S., Duan, S., Allali-Hassani, A., Zuo, X., et al. (2011). Recognition of multivalent histone states associated with heterochromatin by UHRF1 protein. J. Biol. Chem. 286, 24300–24311.

70. Nagymihály, M., Veluchamy, A., Györgypál, Z., Ariel, F., Jégu, T., Benhamed, M., Szücs, A., Kereszt, A., Mergaert, P., and Kondorosi, É. (2017). Ploidy-dependent changes in the epigenome of symbiotic cells correlate with specific patterns of gene expression. Proc. Natl. Acad. Sci. U. S. A. 114, 4543–4548.

71. Naydenov, M., Baev, V., Apostolova, E., Gospodinova, N., Sablok, G., Gozmanova, M., and Yahubyan, G. (2015). High-temperature effect on genes engaged in DNA methylation and affected by DNA methylation in Arabidopsis. Plant Physiol Biochem 87, 102–108.

72. Nishiyama, A., Yamaguchi, L., Sharif, J., Johmura, Y., Kawamura, T., Nakanishi, K., Shimamura, S., Arita, K., Kodama, T., Ishikawa, F., et al. (2013). Uhrf1-dependent H3K23 ubiquitylation couples maintenance DNA methylation and replication. Nature 502, 249–253.

73. Pajoro, A., Severing, E., Angenent, G.C., and Immink, R.G.H. (2017). Histone H3 lysine 36 methylation affects temperature-induced alternative splicing and flowering in plants. Genome Biol. 18, 102.

74. Parent, B., and Tardieu, F. (2012). Temperature responses of developmental processes have not been affected by breeding in different ecological areas for 17 crop species. New Phytol. 194, 760– 774.

75. Peña, P. V., Davrazou, F., Shi, X., Walter, K.L., Verkhusha, V. V., Gozani, O., Zhao, R., and Kutateladze, T.G. (2006). Molecular mechanism of histone H3K4me3 recognition by plant homeodomain of ING2. Nature 442, 100–103.

76. Piacentini, L., Fanti, L., Negri, R., Del Vescovo, V., Fatica, A., Altieri, F., and Pimpinelli, S. (2009). Heterochromatin Protein 1 (HP1a) positively regulates euchromatic gene expression through RNA transcript association and interaction with hnRNPs in Drosophila. PLoS Genet. 5, e1000670.

77. Pikaard, C.S., Haag, J.R., Ream, T., and Wierzbicki, A.T. (2008). Roles of RNA polymerase IV in gene silencing. Trends Plant Sci. 13, 390–397.

78. Popova, O. V, Dinh, H.Q., Aufsatz, W., and Jonak, C. (2013). The RdDM Pathway Is Required for Basal Heat Tolerance in Arabidopsis. Mol. Plant 6, 396–410.

79. Qian, X., Zhao, J., Yeung, P.Y., Zhang, Q.C., and Kwok, C.K. (2019). Revealing lncRNA Structures and Interactions by Sequencing-Based Approaches. Trends Biochem. Sci. 44, 33–52.

80. Ramsahoye, B.H., Biniszkiewicz, D., Lyko, F., Clark, V., Bird, A.P., and Jaenisch, R. (2000). Non-CpG methylation is prevalent in embryonic stem cells and may be mediated by DNA methyltransferase 3a. Proc. Natl. Acad. Sci. U. S. A. 97, 5237–5242.

81. Rigo, R., Bazin, J., Romero-Barrios, N., Moison, M., Lucero, L., Christ, A., Benhamed, M., Blein, T., Huguet, S., Charon, C., et al. (2020). The Arabidopsis lncRNA ASCO modulates the transcriptome through interaction with splicing factors. EMBO Rep. 21, e48977.

82. Rizzardi, K., Landberg, K., Nilsson, L., Ljung, K., and Sundås-Larsson, A. (2011). TFL2/LHP1 is involved in auxin biosynthesis through positive regulation of YUCCA genes. Plant J. 65, 897–906.

83. Rose, N.R., and Klose, R.J. (2014). Understanding the relationship between DNA methylation and histone lysine methylation. Biochim. Biophys. Acta - Gene Regul. Mech. 1839, 1362–1372.

84. Rothbart, S.B., Krajewski, K., Nady, N., Tempel, W., Xue, S., Badeaux, A.I., Barsyte-Lovejoy, D., Martinez, J.Y., Bedford, M.T., Fuchs, S.M., et al. (2012). Association of UHRF1 with methylated H3K9 directs the maintenance of DNA methylation. Nat. Struct. Mol. Biol. 19, 1155–1160.

85. Di Ruscio, A., Ebralidze, A.K., Benoukraf, T., Amabile, G., Goff, L.A., Terragni, J., Figueroa, M.E., De Figueiredo Pontes, L.L., Alberich-Jorda, M., Zhang, P., et al. (2013). DNMT1-interacting RNAs block gene-specific DNA methylation. Nature 503, 371–376.

86. Sakata, T., Oshino, T., Miura, S., Tomabechi, M., Tsunaga, Y., Higashitani, N., Miyazawa, Y., Takahashi, H., Watanabe, M., and Higashitani, A. (2010). Auxins reverse plant male sterility caused by high temperatures. Proc. Natl. Acad. Sci. U. S. A. 107, 8569–8574.

87. Schindelin, J., Arganda-Carreras, I., Frise, E., Kaynig, V., Longair, M., Pietzsch, T., Preibisch, S., Rueden, C., Saalfeld, S., Schmid, B., et al. (2012). Fiji: An open-source platform for biological-image analysis. Nat. Methods 9, 676–682.

88. Severing, E., Faino, L., Jamge, S., Busscher, M., Kuijer-Zhang, Y., Bellinazzo, F., Busscher-Lange, J., Fernández, V., Angenent, G.C., Immink, R.G.H., et al. (2018). Arabidopsis thaliana ambient temperature responsive lncRNAs. BMC Plant Biol. 18, 145.

89. Sharif, J., Muto, M., Takebayashi, S.I., Suetake, I., Iwamatsu, A., Endo, T.A., Shinga, J., Mizutani-Koseki, Y., Toyoda, T., Okamura, K., et al. (2007). The SRA protein Np95 mediates epigenetic inheritance by recruiting Dnmt1 to methylated DNA. Nature 450, 908–912.

90. Shen, X., De Jonge, J., Forsberg, S.K.G., Pettersson, M.E., Sheng, Z., Hennig, L., and Carlborg, Ö. (2014). Natural CMT2 Variation Is Associated With Genome-Wide Methylation Changes and Temperature Seasonality. PLoS Genet. 10, e1004842.

91. Shi, X., Hong, T., Walter, K.L., Ewalt, M., Michishita, E., Hung, T., Carney, D., Peña, P., Lan, F., Kaadige, M.R., et al. (2006). ING2 PHD domain links histone H3 lysine 4 methylation to active gene repression. Nature 442, 96–99.

92. Shook, M.S., and Richards, E.J. (2014). VIM proteins regulate transcription exclusively through the MET1 cytosine methylation pathway. Epigenetics 9, 980–986.

93. Sidaway-Lee, K., Costa, M.J., Rand, D.A., Finkenstadt, B., and Penfield, S. (2014). Direct measurement of transcription rates reveals multiple mechanisms for configuration of the Arabidopsis ambient temperature response. Genome Biol. 15, R45.

94. Siegfried, N.A., Busan, S., Rice, G.M., Nelson, J.A.E., and Weeks, K.M. (2014). RNA motif discovery by SHAPE and mutational profiling (SHAPE-MaP). Nat. Methods 11, 959–965.

95. Skourti-Stathaki, K., Torlai Triglia, E., Warburton, M., Voigt, P., Bird, A., and Pombo, A. (2019). R-Loops Enhance Polycomb Repression at a Subset of Developmental Regulator Genes. Mol. Cell 73, 930–945.e4.

96. Smallwood, A., Estève, P.O., Pradhan, S., and Carey, M. (2007). Functional cooperation between HP1 and DNMT1 mediates gene silencing. Genes Dev. 21, 1169–1178.

97. Smola, M.J., and Weeks, K.M. (2018). In-cell RNA structure probing with SHAPE-MaP. Nat. Protoc. 13, 1181–1195.

98. Somero, G.N. (2018). RNA thermosensors: How might animals exploit their regulatory potential? J. Exp. Biol. 221.

99. Soppe, W.J.J., Jasencakova, Z., Houben, A., Kakutani, T., Meister, A., Huang, M.S., Jacobsen, S.E., Schubert, I., and Fransz, P.F. (2002). DNA methylation controls histone H3 lysine 9 methylation and heterochromatin assembly in Arabidopsis. EMBO J. 21, 6549–6559.

100. Sorenson, R., and Bailey-Serres, J. (2015). Rapid immunopurification of ribonucleoprotein complexes of plants. Methods Mol. Biol. 1284, 209–219.

101. Spitale, R.C., Flynn, R.A., Zhang, Q.C., Crisalli, P., Lee, B., Jung, J.W., Kuchelmeister, H.Y., Batista, P.J., Torre, E.A., Kool, E.T., et al. (2015). Structural imprints in vivo decode RNA regulatory mechanisms. Nature 519, 486–490.

102. Steffen, A., and Staiger, D. (2017). Chromatin marks and ambient temperature-dependent flowering strike up a novel liaison. Genome Biol. 18, 119.

103. Stroud, H., Greenberg, M.V.C., Feng, S., Bernatavichute, Y. V., and Jacobsen, S.E. (2013). Comprehensive analysis of silencing mutants reveals complex regulation of the Arabidopsis methylome. Cell 152, 352–364.

104. Stroud, H., Do, T., Du, J., Zhong, X., Feng, S., Johnson, L., Patel, D.J., and Jacobsen, S.E. (2014). Non-CG methylation patterns shape the epigenetic landscape in Arabidopsis. Nat. Struct. Mol. Biol. 21, 64–72.

105. Susila, H., Nasim, Z., and Ahn, J.H. (2018). Ambient temperature-responsive mechanisms coordinate regulation of flowering time. Int. J. Mol. Sci. 19, 3196.

106. Suzuki, M.M., and Bird, A. (2008). DNA methylation landscapes: Provocative insights from epigenomics. Nat. Rev. Genet. 9, 465–476.

107. Tamaru, H., and Selker, E.U. (2001). A histone H3 methyltransferase controls DNA methylation in Neurospora crassa. Nature 414, 277–283.

108. Taniue, K., Kurimoto, A., Sugimasa, H., Nasu, E., Takeda, Y., Iwasaki, K., Nagashima, T., Okada-Hatakeyama, M., Oyama, M., Kozuka-Hata, H., et al. (2016). Long noncoding RNA UPAT promotes colon tumorigenesis by inhibiting degradation of UHRF1. Proc. Natl. Acad. Sci. U. S. A. 113, 1273– 1278.

109. Tariq, M., Saze, H., Probst, A. V., Lichota, J., Habu, Y., and Paszkowski, J. (2003). Erasure of CpG methylation in Arabidopsis alters patterns of histone H3 methylation in heterochromatin. Proc. Natl. Acad. Sci. U. S. A. 100, 8823–8827.

110. Tasset, C., Singh Yadav, A., Sureshkumar, S., Singh, R., van der Woude, L., Nekrasov, M., Tremethick, D., van Zanten, M., and Balasubramanian, S. (2018). POWERDRESS-mediated histone deacetylation is essential for thermomorphogenesis in Arabidopsis thaliana. PLOS Genet. 14, e1007280.

111. Turck, F., Roudier, F., Farrona, S., Martin-Magniette, M.-L., Guillaume, E., Buisine, N., Gagnot, S., Martienssen, R.A., Coupland, G., and Colot, V. (2007). Arabidopsis TFL2/LHP1 Specifically Associates with Genes Marked by Trimethylation of Histone H3 Lysine 27. PLoS Genet. 3, e86.

112. Unoki, M., Nishidate, T., and Nakamura, Y. (2004). ICBP90, an E2F-1 target, recruits HDAC1 and binds to methyl-CpG through its SRA domain. Oncogene 23, 7601–7610.

113. Varley, K.E., Gertz, J., Bowling, K.M., Parker, S.L., Reddy, T.E., Pauli-Behn, F., Cross, M.K., Williams, B.A., Stamatoyannopoulos, J.A., Crawford, G.E., et al. (2013). Dynamic DNA methylation across diverse human cell lines and tissues. Genome Res. 23, 555–567.

114. Veluchamy, A., Jégu, T., Ariel, F., Latrasse, D., Mariappan, K.G., Kim, S.K., Crespi, M., Hirt, H., Bergounioux, C., Raynaud, C., et al. (2016). LHP1 Regulates H3K27me3 Spreading and Shapes the Three-Dimensional Conformation of the Arabidopsis Genome. PLoS One 11, e0158936.

115. Vernié, T., Kim, J., Frances, L., Ding, Y., Sun, J., Guan, D., Niebel, A., Gifford, M.L., de Carvalho-Niebel, F., and Oldroyd, G.E.D. (2015). The NIN Transcription Factor Coordinates Diverse Nodulation Programs in Different Tissues of the Medicago truncatula Root. Plant Cell 27, 3410– 3424.

116. Waadt, R., and Kudla, J. (2008). In planta visualization of protein interactions using bimolecular fluorescence complementation (BiFC). Cold Spring Harb. Protoc. 3.

117. Wang, H., Cao, D., and Wu, F. (2018). Long noncoding RNA UPAT promoted cell proliferation via increasing UHRF1 expression in non-small cell lung cancer. Oncol. Lett. 16, 1491–1498.

118. Wang, Y., Fan, X., Lin, F., He, G., Terzaghi, W., Zhu, D., and Deng, X.W. (2014). Arabidopsis noncoding RNA mediates control of photomorphogenesis by red light. Proc. Natl. Acad. Sci. U. S. A. 111, 10359–10364.

119. Wassenegger, M., Heimes, S., Riedel, L., and Sänger, H.L. (1994). RNA-directed de novo methylation of genomic sequences in plants. Cell 76, 567–576.

120. Winter, D., Vinegar, B., Nahal, H., Ammar, R., Wilson, G. V., and Provart, N.J. (2007). An “Electronic Fluorescent Pictograph” Browser for Exploring and Analyzing Large-Scale Biological Data Sets. PLoS One 2, e718.

121. Woo, H.R., Pontes, O., Pikaard, C.S., and Richards, E.J. (2007). VIM1, a methylcytosine-binding protein required for centromeric heterochromatinization. Genes Dev. 21, 267–277.

122. Woo, H.R., Dittmer, T.A., and Richards, E.J. (2008). Three SRA-Domain Methylcytosine-Binding Proteins Cooperate to Maintain Global CpG Methylation and Epigenetic Silencing in Arabidopsis. PLoS Genet. 4.

123. Wysocka, J., Swigut, T., Xiao, H., Milne, T.A., Kwon, S.Y., Landry, J., Kauer, M., Tackett, A.J., Chait, B.T., Badenhorst, P., et al. (2006). A PHD finger of NURF couples histone H3 lysine 4 trimethylation with chromatin remodelling. Nature 442, 86–90.

124. Xu, W., Xu, H., Li, K., Fan, Y., Liu, Y., Yang, X., and Sun, Q. (2017). The R-loop is a common chromatin feature of the Arabidopsis genome. Nat. Plants 3, 704–714.

125. Xue, B., Zhao, J., Feng, P., Xing, J., Wu, H., and Li, Y. (2019). Epigenetic mechanism and target therapy of uhrf1 protein complex in malignancies. Onco. Targets. Ther. 12, 549–559.

126. Yao, Y., Bilichak, A., Golubov, A., and Kovalchuk, I. (2012). ddm1 plants are sensitive to methyl methane sulfonate and NaCl stresses and are deficient in DNA repair. Plant Cell Rep. 31, 1549– 1561.

127. Yu, M., and Ren, B. (2017). The Three-Dimensional Organization of Mammalian Genomes. Annu. Rev. Cell Dev. Biol. 33, 265–289.

128. Yu, Y., Dong, A., and Shen, W.H. (2004). Molecular characterization of the tobacco SET domain protein NtSET1 unravels its role in histone methylation, chromatin binding, and segregation. Plant J. 40, 699–711.

129. Zemach, A., Kim, M.Y., Hsieh, P.H., Coleman-Derr, D., Eshed-Williams, L., Thao, K., Harmer, S.L., and Zilberman, D. (2013). The nucleosome remodeler DDM1 allows DNA methyltransferases to access H1-containing heterochromatin. Cell 153, 193–205.

130. Zhang, X., Yazaki, J., Sundaresan, A., Cokus, S., Chan, S.W.L., Chen, H., Henderson, I.R., Shinn, P., Pellegrini, M., Jacobsen, S.E., et al. (2006). Genome-wide High-Resolution Mapping and Functional Analysis of DNA Methylation in Arabidopsis. Cell 126, 1189–1201.

131. Zhang, Y., Chen, M., Siemiatkowska, B., Toleco, M.R., Jing, Y., Strotmann, V., Zhang, J., Stahl, Y., and Fernie, A.R. (2020). A Highly Efficient Agrobacterium-Mediated Method for Transient Gene Expression and Functional Studies in Multiple Plant Species. Plant Commun. 1, 100028.

132. Zhao, Q., Zhang, J., Chen, R., Wang, L., Li, B., Cheng, H., Duan, X., Zhu, H., Wei, W., Li, J., et al. (2016). Dissecting the precise role of H3K9 methylation in crosstalk with DNA maintenance methylation in mammals. Nat. Commun. 7, 1–12.

