## Supplemental Figures 1 to 11 and Tables 1 and 4 for "Sequence-unrelated long noncoding RNAs converged to modulate the activity of conserved epigenetic machineries across kingdoms"

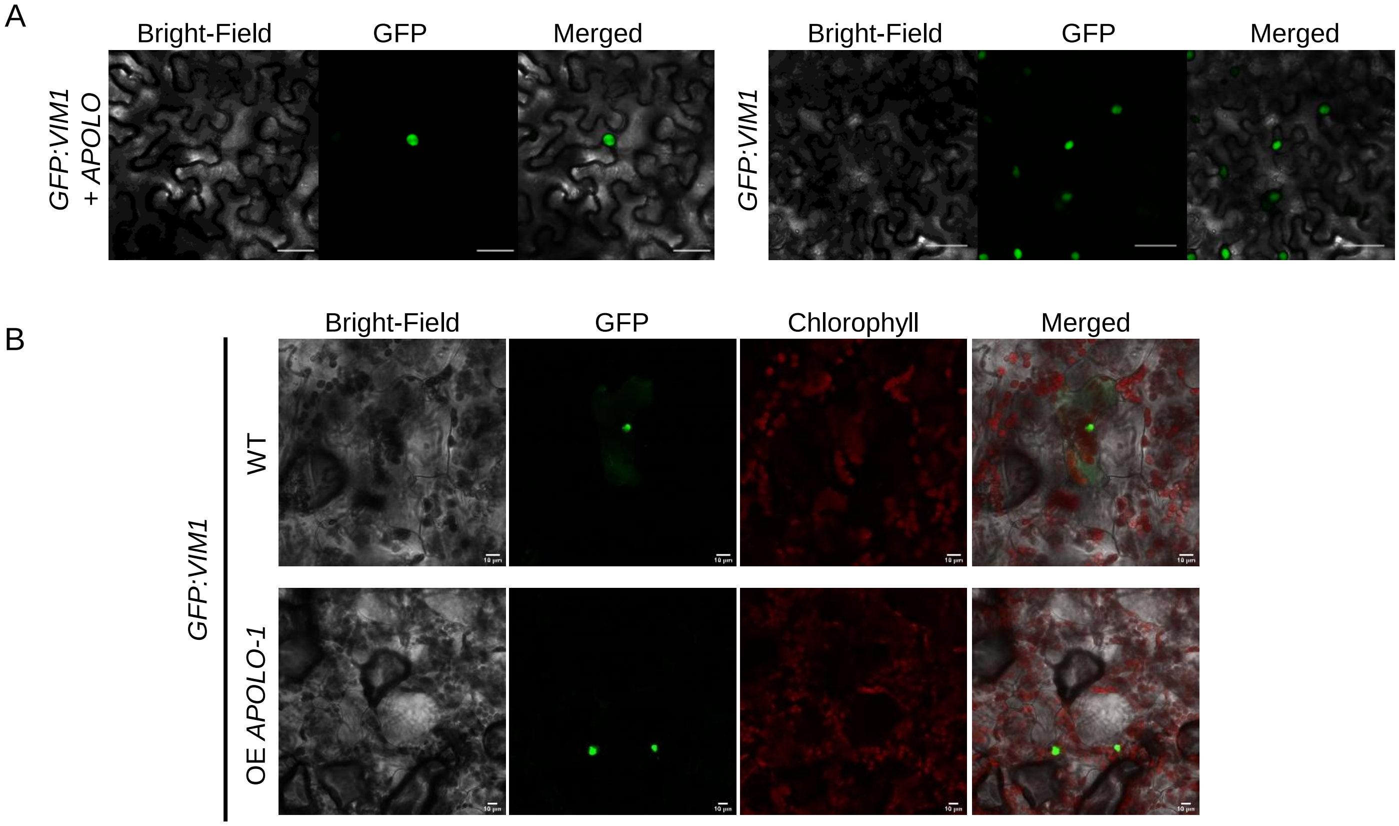


**Supplementary Figure 1: GFP-VIM1 localizes at the nucleus**

(**A**) Sub-cellular localization of GFP-VIM1 translational fusion transiently expressed under the control of the 35S-CaMV promoter in *Nicotiana benthamiana* leaves in presence (left panel) or absence (right panel) of *APOLO*. Bright-field image (left), GFP fluorescence alone (middle) and bright-field merged images (right) are shown. Scale bars, 50µm. (**B**) Sub-cellular localization of GFP-VIM1 translational fusion transiently expressed under the control of the 35S-CaMV promoter in *Arabidopsis thaliana* wild-type (WT) and *APOLO* over-expression (OE *APOLO-1*) leaves. Bright-field image (left), GFP or chlorophyll fluorescence alone (middle), and bright-field merged images (right) are shown. Scale bars, 10µm.
In (**A**) and (**B**), one representative picture out of three biological replicates is shown.

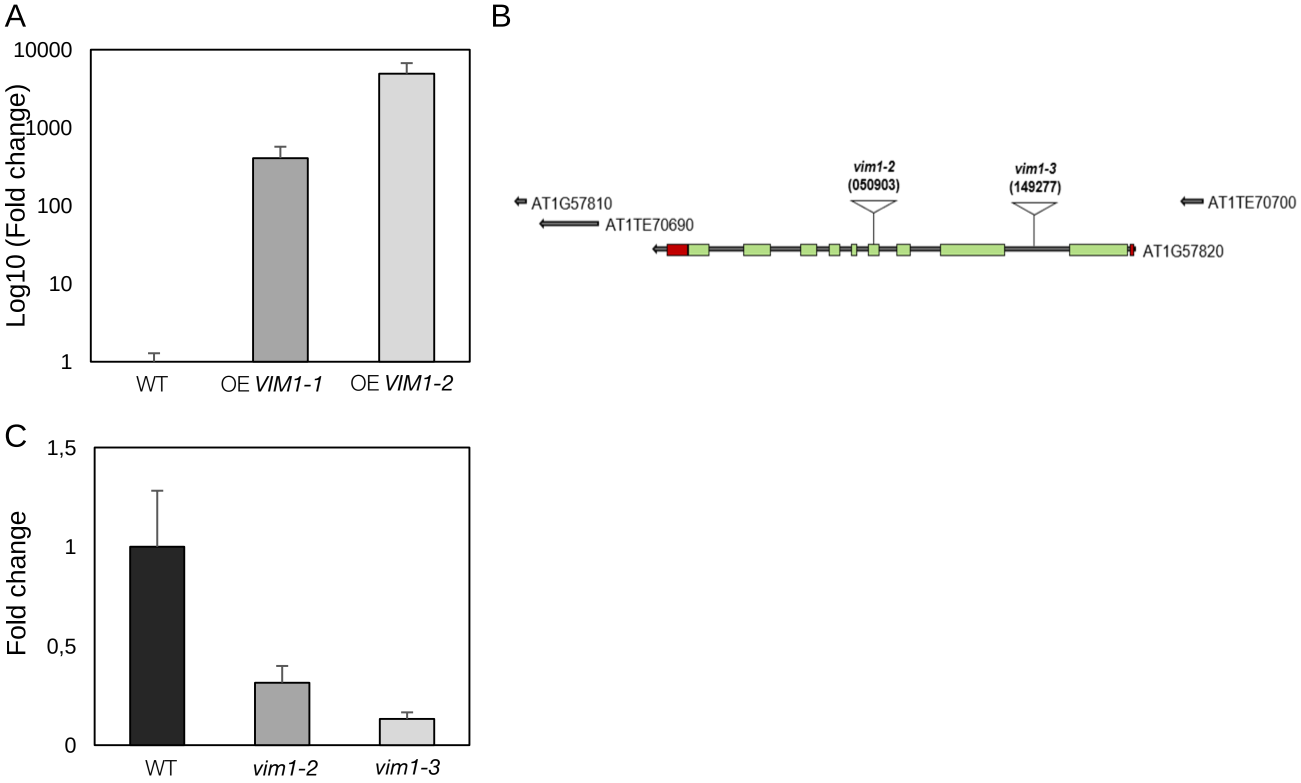


**Supplementary Figure 2: Characterization of *Arabidopsis thaliana VIM1* over-expression and *vim1* homozygous T-DNA insertion lines**

(**A**) *VIM1* basal transcript levels in wild-type (WT), *VIM1-1* (OE *VIM1-1*) and *VIM1-2* (OE *VIM1-2*) over-expression lines. (**B**) Schematic representation of the intron-exon structure of the *VIM1* gene (AT1G57820) and positions of T-DNA insertions with allele designations (SALK collection: *vim1-2*: 050903; *vim1-3*: 149277). Exons are indicated by green boxes and introns by grey lines. 5' and 3' UTR regions are indicated by red boxes. AT1G57810 (reverse transcriptase pseudogene), AT1TE70690 and AT1TE70700 (LINE1 transposons) located near AT1G57820 are indicated by grey arrows. (**C**) *VIM1* basal transcript levels in WT, *vim1-2* and *vim1-3* homozygous T-DNA insertion lines.
In (**A**), (**C**), transcript levels are normalized to *PP2A* expression levels. Bars represent average ± SD (n = 3 independent pools of seedlings).

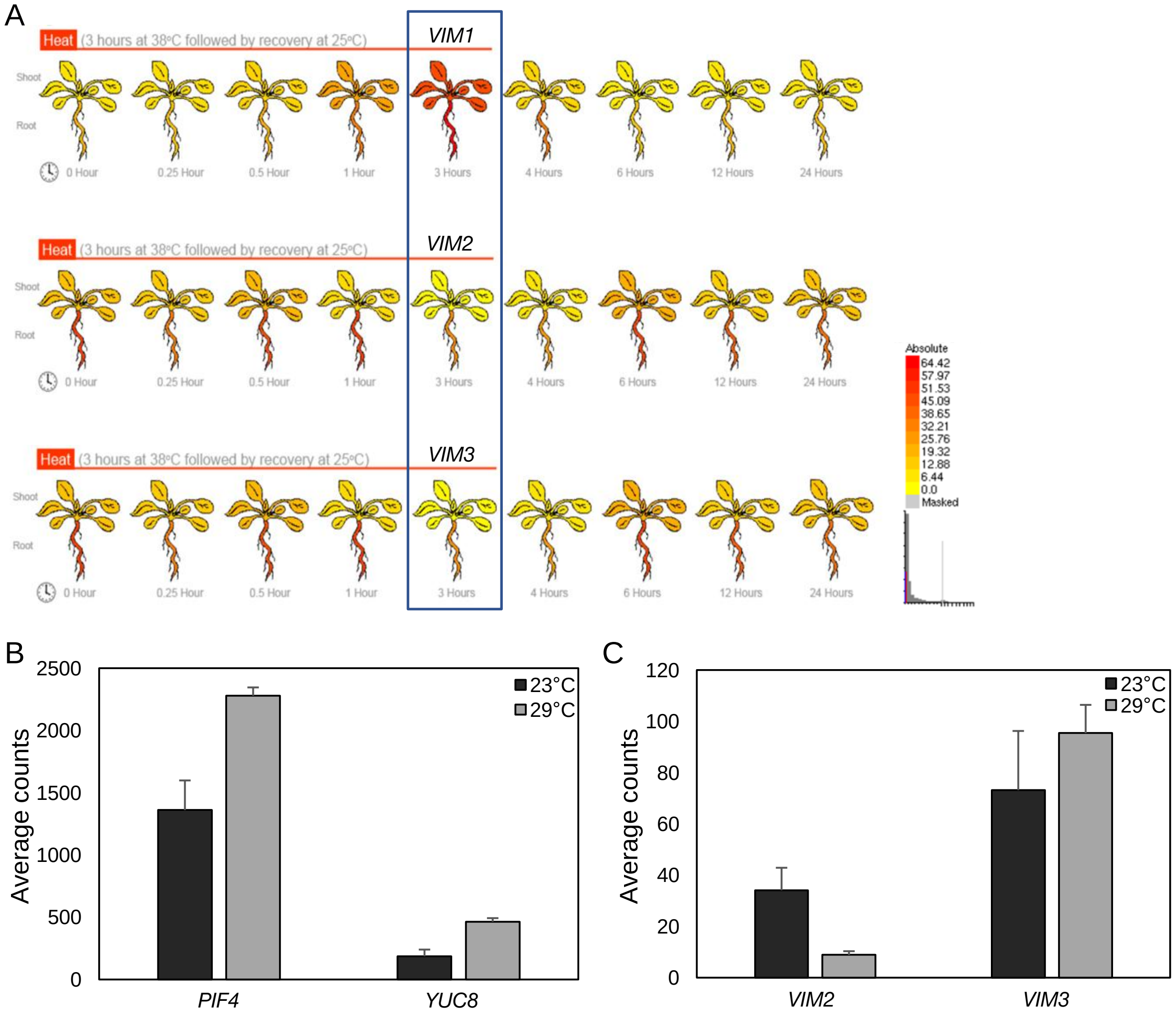


**Supplementary Figure 3: *VIM1*, *VIM2* and *VIM3* are differentially regulated in reponse to heat**

(**A**) eFP browser view of *VIM1*, *VIM2* and *VIM3* expression patterns in Arabidopsis shoot and root in response to heat (3 hours at 38°C followed by recovery at 25°C). Expression strength coded by color: yellow = low, red = high. The Arabidopsis eFP Browser is located at bar.utoronto.ca and published in Winter et al. (2007). The blue rectangle points at the major difference among genes. (**B-C**) Expression level of the heat markers *PIF4* and *YUC8* (in **B**), and *VIM1* homologs *VIM2* and *VIM3* (in **C**), in 4-day-old wild-type (WT) seedlings treated with heat (29°C) for 6h, related to Figure 1.

In (**B-C**), results are expressed as normalized counts obtained by RNA-sequencing. Bars represent average ± SD (n = 3 biological replicates).

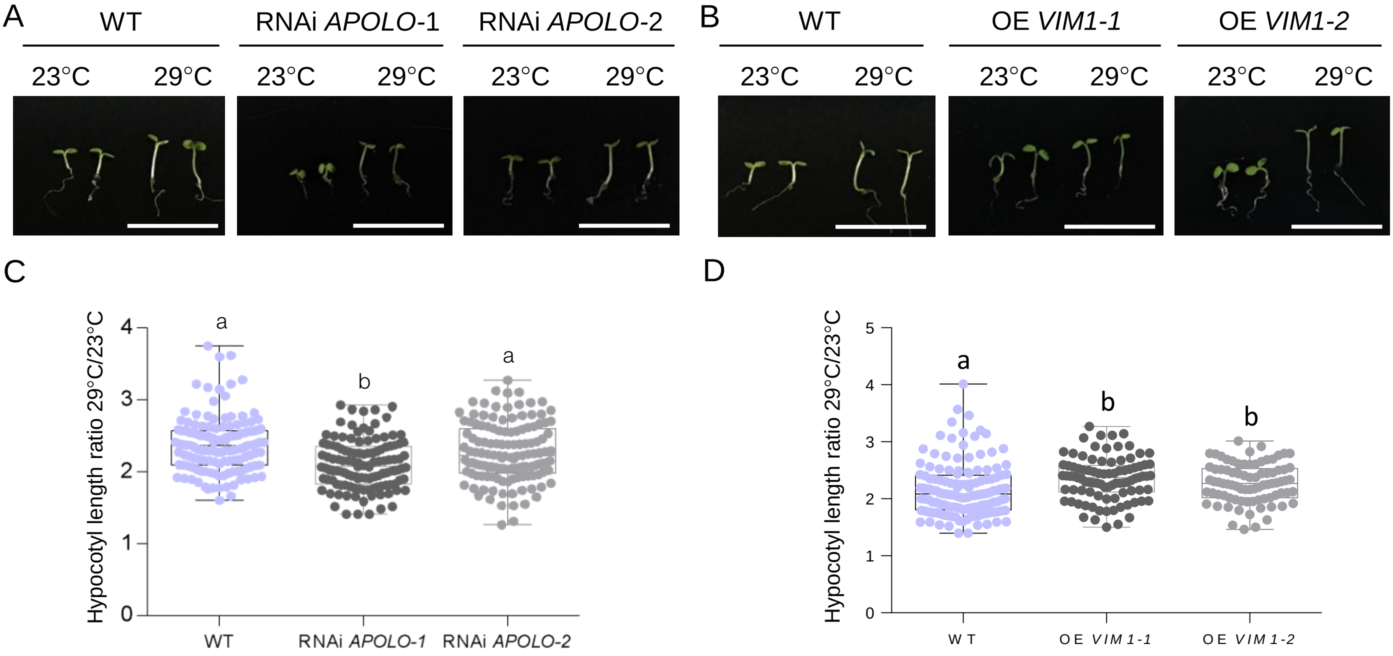


**Supplementary Figure 4: *APOLO* and VIM1 regulate hypocotyle elongation in response to heat**

(A-B) Representative morphological phenotypes of 4-day-old RNAi *APOLO* (RNAi *APOLO*-1, RNAi *APOLO*-2) (A) or *VIM1* overexpression (OE *VIM1-1*, OE *VIM1-2*) (B) seedlings and their associated wild-type (WT) grown at 23°C or 29°C. Scale bars, 1 cm. (C-D) Boxplots showing hypocotyl length at 29°C over 23°C of 4-day-old RNAi *APOLO* (C) or *VIM1* over-expression (D) seedlings and their associated WT. Values are represented by colored points.

In (C-D), results are the mean of three biological replicates and letters indicate significant differences compared to WT, based on a Kruskal-Wallis test ($\alpha = 0.05$; $n \geq 81$).

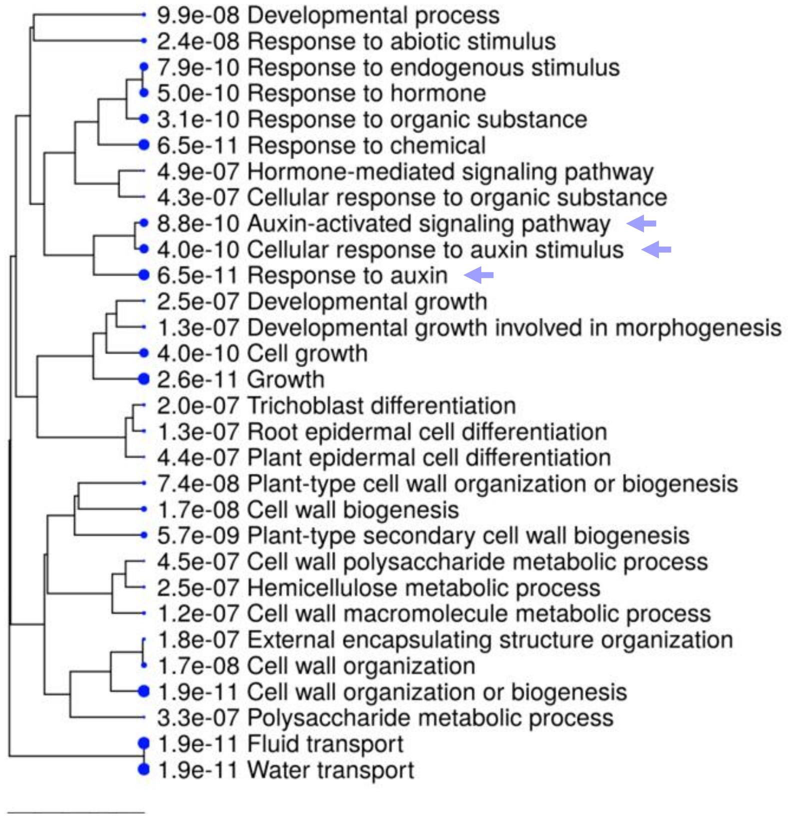


**Supplementary Figure 5: Gene categories transcriptionally regulated in thermomorphogenesis**

Gene ontology (GO) enrichment analyses of upregulated transcripts in 4-day-old wild-type (WT) seedlings treated with heat (29°C) for 6h. The hierarchical clustering trees summarize the correlation among significant pathways. Pathways with many shared genes are clustered together. Bigger dots indicate more significant P-values. Arrows indicate auxin-related pathways. The ShinyGO Browser is located at bioinformatics.sdstate.edu and published in Ge and Jung (2018).

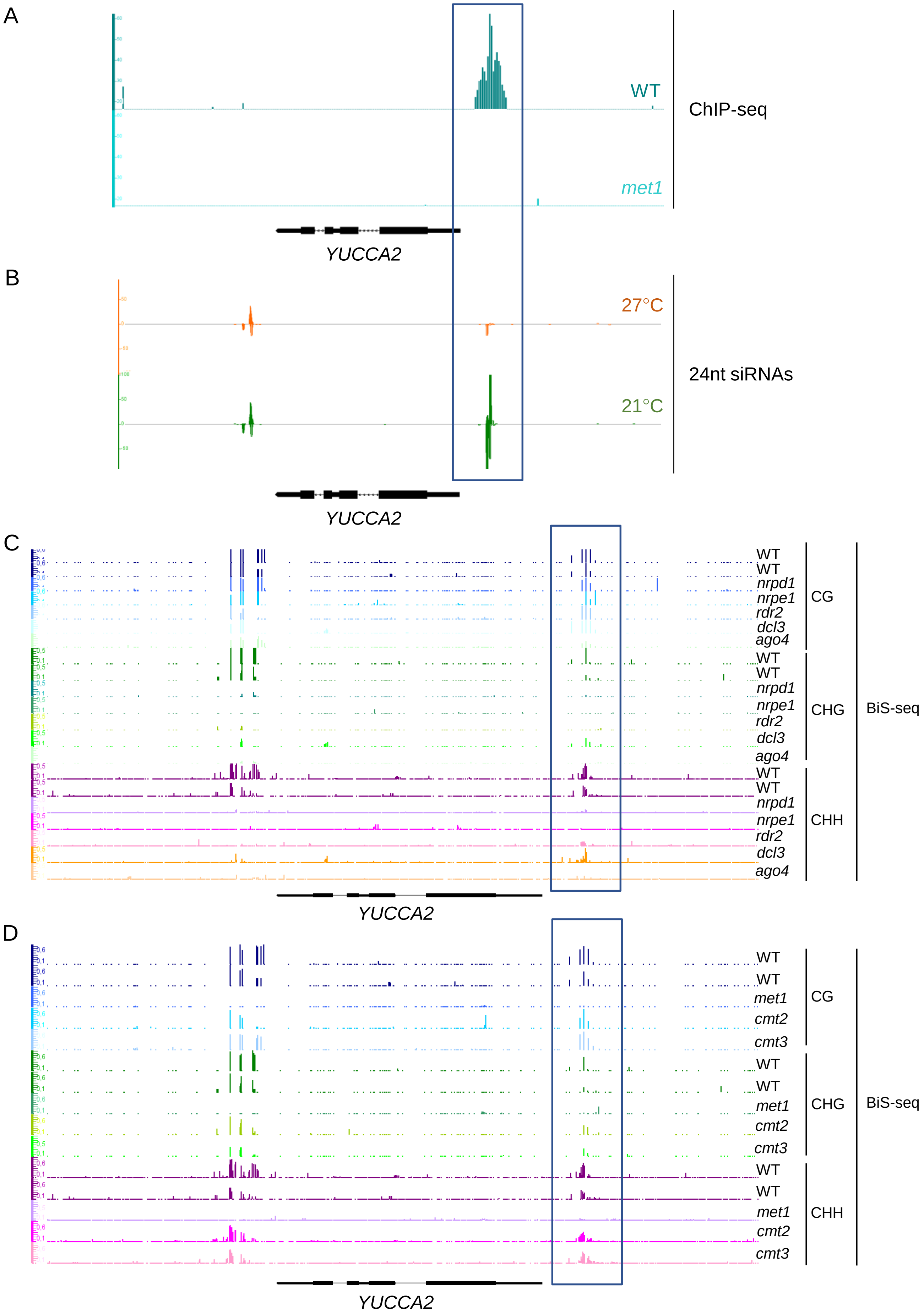


**Supplementary Figure 6: Epigenetic profile of the *YUCCA2* locus**

(**A**) PolV binding at the *YUCCA2* (*YUC2*) locus by NRPE1 subunit-chromatin immunoprecipitation (ChIP)-sequencing in wild-type (WT; Track 1) and *met1* mutant (Track 2) (Johnson et al. 2014). (**B**) 24nt siRNA coverage at the *YUC2* locus at 27°C (Track 1) or 21°C (Track 2) (Gyula et al. 2018). (**C**) Distribution of DNA methylation at the *YUC2* locus by bisulfite (BiS)-sequencing, in the three sequence contexts CG (Tracks 1 to 7), CHG (Tracks 8 to 14) and CHH (Tracks 15 to 21), in *nrpd1* (Tracks 3, 10, 17), *nrpe1* (Tracks 4, 11, 18), *rdr2* (Tracks 5, 12, 19), *ago4* (Tracks 6, 13, 20) or *dcl3* (Tracks 7, 14, 21) mutants and their associated WT (Tracks 1, 2, 8, 9, 15, 16) (Stroud et al. 2013). (**D**) Distribution of DNA methylation at the *YUC2* locus by BiS-seq, in the three sequence contexts CG (Tracks 1 to 5), CHG (Tracks 6 to 10) and CHH (Tracks 11 to 15), in *met1* (Tracks 3, 8, 13), *cmt2* (Tracks 4, 9, 14), or *cmt3* (Tracks 5, 10, 15) mutants and their associated WT (Tracks 1, 2, 6, 7, 11, 12) (Stroud et al. 2013).
In (**A-D**), gene annotation is shown at the bottom. The vertical rectangles highlight the same region of interest in all panels.

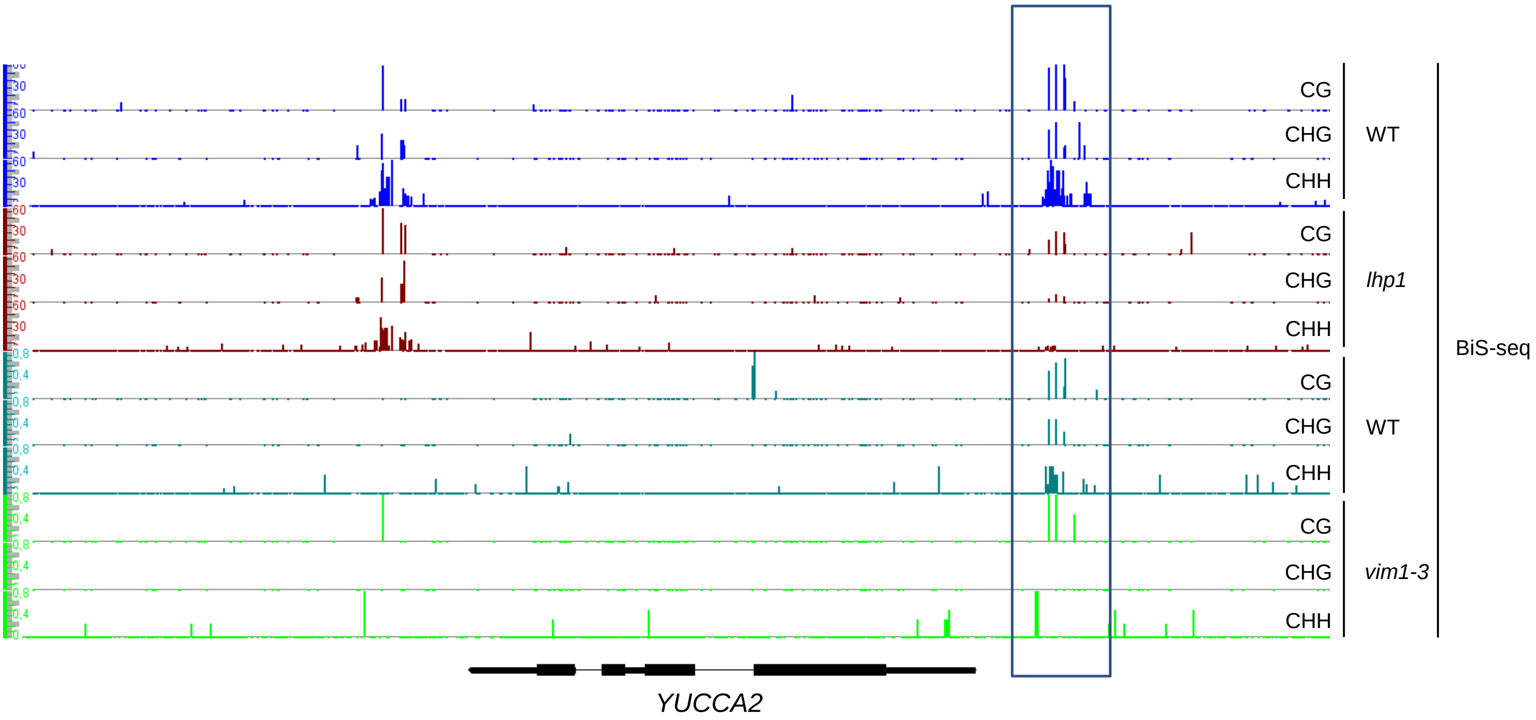


**Supplementary Figure 7: VIM1 and LHP1 regulate methylation at the promoter of the thermomorphogenesis-related gene *YUCCA2***

Distribution of DNA methylation at the *YUCCA2* (*YUC2*) locus by bisulfite (BiS)-sequencing, in the three sequence contexts CG (Tracks 1, 4, 7, 10), CHG (Tracks 2, 5, 8, 11) and CHH (Tracks 3, 6, 9, 12), in *lhp1* (Tracks 4 to 6) or *vim1-3* (Tracks 10 to12) mutant seedlings and their associated WT (Tracks 1 to 3 and 7 to 9, respectively). Gene annotation is shown in the bottom. The vertical rectangle highlights the same region of interest as in Supplementary Figure 6.

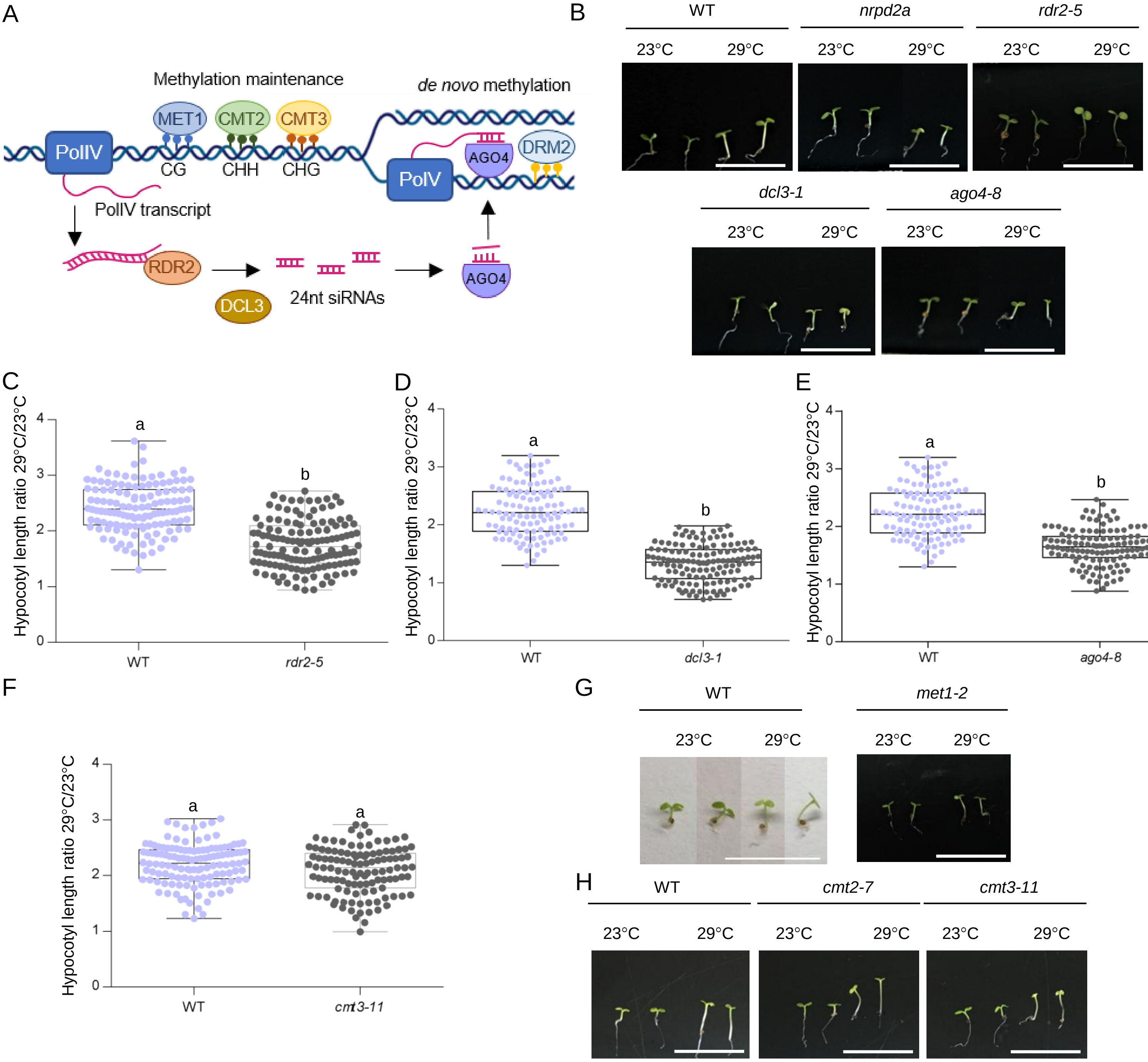


**Supplementary Figure 8: *de novo* DNA methylation and its maintenance regulate thermomorphogenesis**

(**A**) Schematic representation of maintenance and *de novo* DNA methylationthe induced by the RNA-directed DNA methylation (RdDM) pathway. AGO4: ARGONAUTE 4; CMT2: CHROMOMETHYLASE 2; CMT3: CHROMOMETHYLASE 3; DCL3: DICER-LIKE 3; DRM2: DOMAINS REARRANGED METHYLTRANSFERASE 2; MET1: METHYLTRANSFERASE 1; PolIV: RNA Polymerase IV; PolV: RNA Polymerase V; RDR2: RNA-DEPENDENT RNA POLYMERASE 2; 24nt siRNA: 24 nucleotide small interfering RNA. CG, CHG and CHH represent the three sequence contexts of DNA methylation. H is A, T, or C. (**B**) Representative morphological phenotypes of 4-day-old RdDM-related mutant seedlings *rdr2-5*, *ago4-8*, *dcl3-1* and their associated wild-type (WT). (**C-F**) Boxplots showing hypocotyl length at 29°C over 23°C of 4-day-old RdDM-related mutant seedlings *rdr2-5* (**C**), *dcl3-1* (**D**), *ago4-8* (**E**), methyltransferase mutant seedlings *cmt3-11* (**F**) and their associated wild-type (WT). (**G-I**) Representative morphological phenotypes of 4-day-old methyltransferase mutant *met1-2* (**G**) or *cmt2-7* and *cmt3-11* (**H**) seedlings and their associated WT, grown at 23°C or 29°C. Scale bars, 1 cm.

In (**C-F**), results are the mean of three biological replicates and letters indicate significant differences compared to WT, based on a Kruskal-Wallis test ($\alpha = 0.05$; $n \geq 113$).

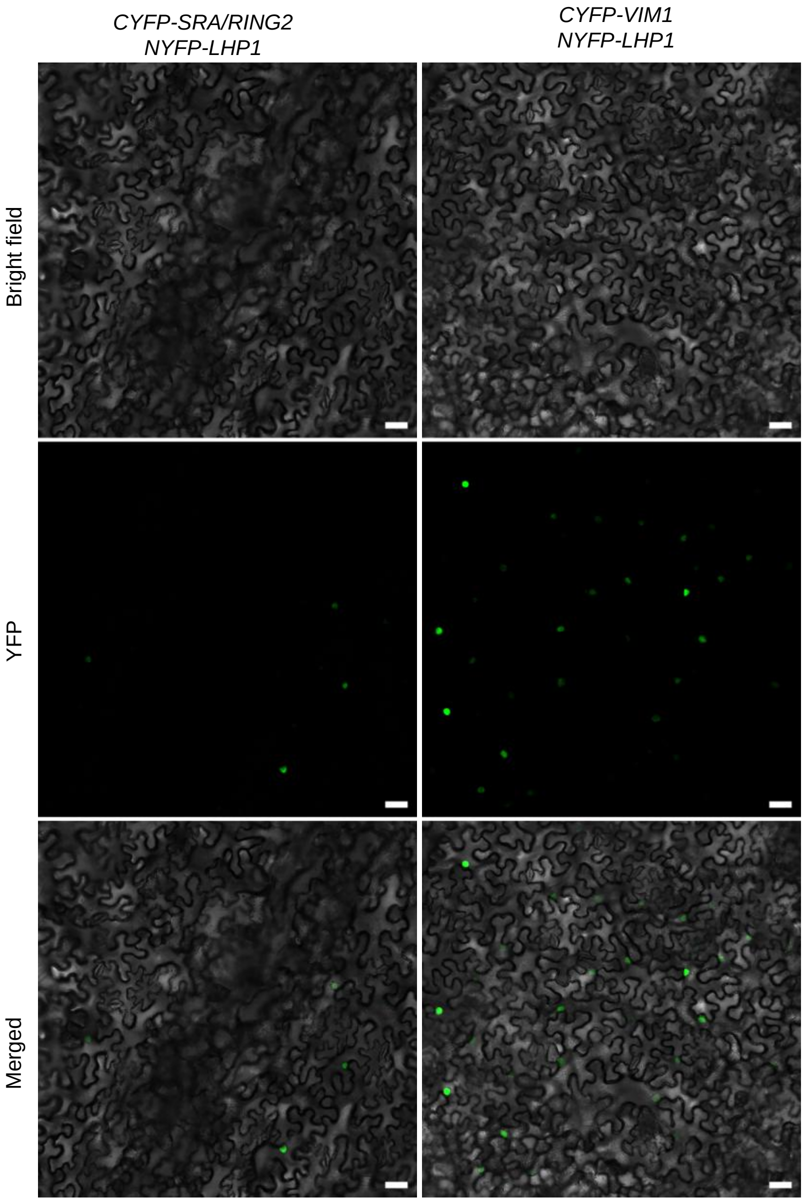


**Supplementary Figure 9: Bimolecular Fluorescence Complementation (BiFC) assay in transiently transformed *Nicotiana benthamiana* leaves**.

CYFP was fused to VIM1 or SRA/RING2 and NYFP was fused to LHP1. Only five nuclei were observed in one out of six biological replicates when CYFP-SRA/RING2 is co-expressed with NYFP-LHP1. In both panels, bright-field images (top), YFP fluorescence alone (middle) and bright-field merged images (bottom) are shown. Scale bars, 50µm.

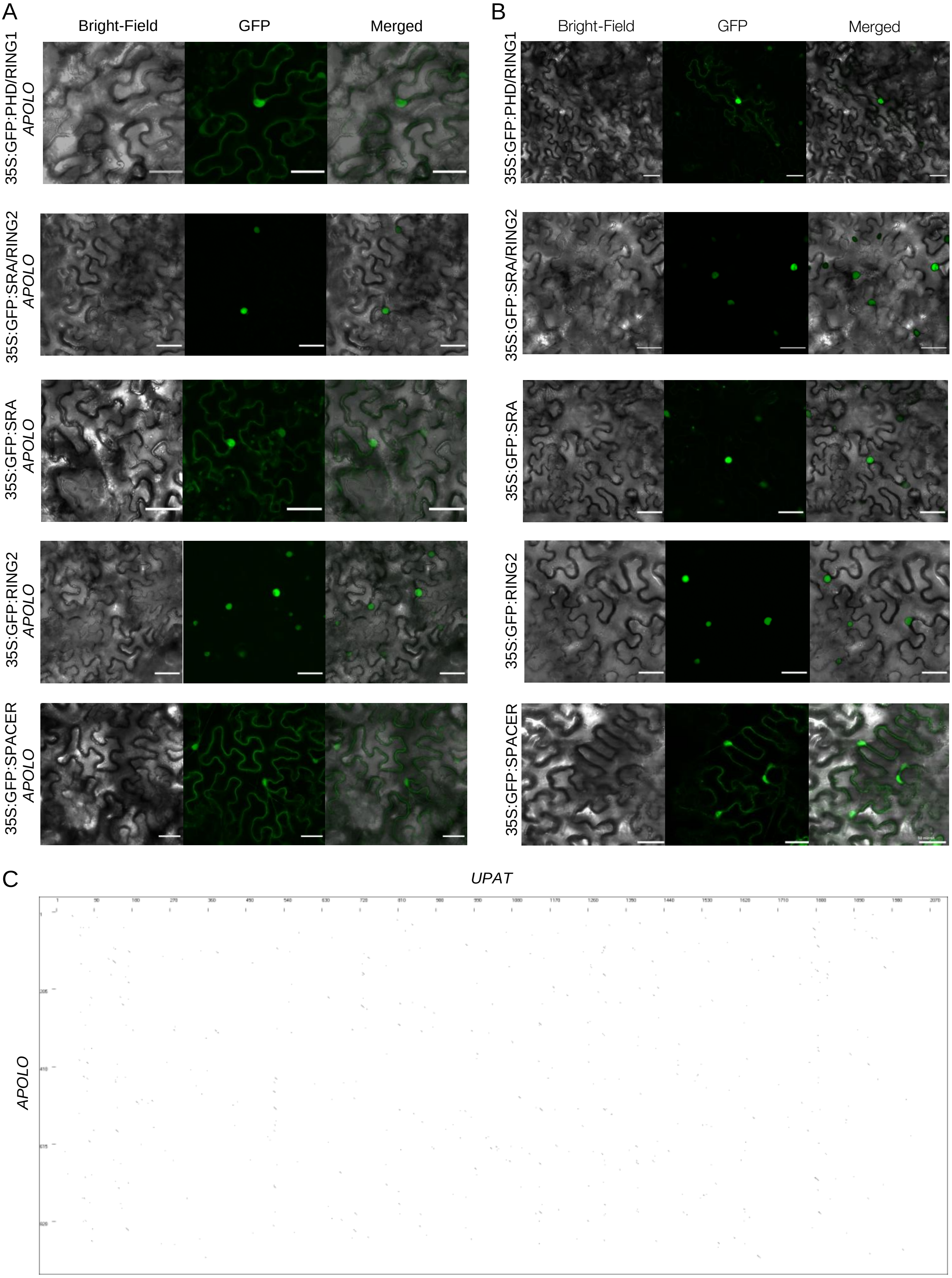


**Supplementary Figure 10: *APOLO* and *UPAT* lncRNAs share mecanisms of interaction with methylcytosine-binding proteins but no sequence similarity**

(**A-B**) Sub-cellular localization of *GFP-PHD/RING1*, *GFP-SRA/RING2*, *GFP-SRA*, *GFP-RING2* and *GFP-SPACER* translational fusions transiently expressed from the 35S-CaMV promoter in *Nicotiana benthamiana* leaves in presence (in **A**) or absence (in **B**) of *APOLO*. Bright-field image (left), GFP fluorescence alone (middle) and bright-field/GFP fluorescence merged images (right) are shown. Scale bars, 50µm. One representative picture out of three biological replicates is shown. (**C**) Dot plot pairwise sequence comparison of *APOLO* and *UPAT* lncRNAs showing low similarity between the two sequences. BioEdit program is published in Hall (1999).

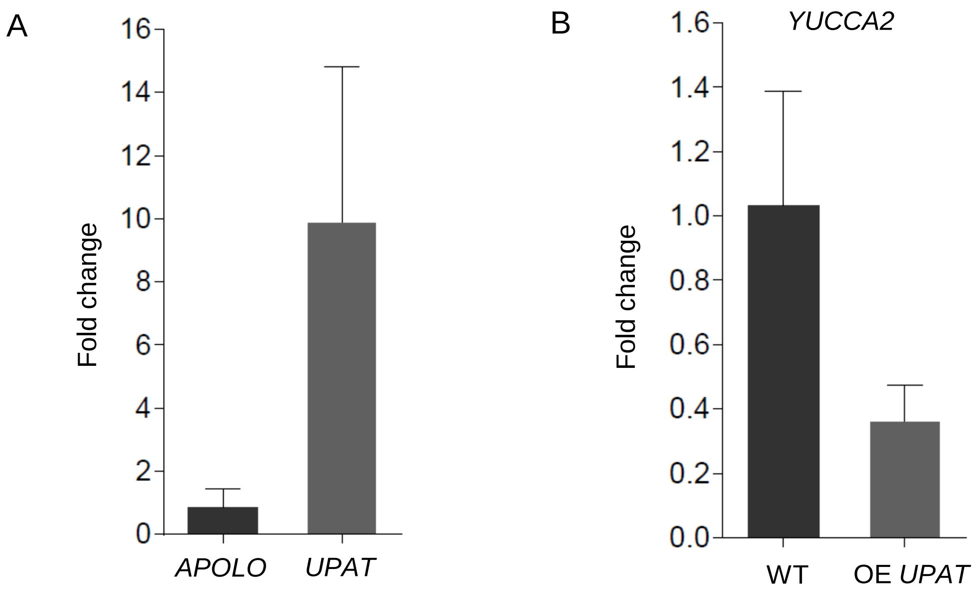


**Supplementary Figure 11: Constitutive expression of the lncRNA *UPAT* in Arabidopsis seedlings impairs *YUCCA2* transcriptional accumulation**

(**A**) *UPAT* transcript levels in OE *UPAT* Arabidopsis seedlings determined as the ratio with endogenous *APOLO* accumulation taken as 1. (**B**) *YUCCA2* basal transcript levels in wild-type (WT) and OE *UPAT* seedlings. Bars represent average ± SD (n = 3 independent pools of seedlings).

| Genotype | Hypocotyl length at 23°C | Hypocotyl length at 29°C |
|---|---|---|
| WT OE *APOLO* | 0,134 ± 0,029 | 0,322 ± 0,077 |
| OE *APOLO-1* | 0,123 ± 0,025 | 0,239 ± 0,052 |
| OE *APOLO-2* | 0,123 ± 0,029 | 0,238 ± 0,056 |
| WT RNAi *APOLO* | 0,134 ± 0,029 | 0,322 ± 0,077 |
| RNAi *APOLO-1* | 0,127 ± 0,029 | 0,267 ± 0,051 |
| RNAi *APOLO-2* | 0,125 ± 0,022 | 0,284 ± 0,057 |
| WT OE *VIM1* | 0,145 ± 0,024 | 0,313 ± 0,066 |
| OE *VIM1-1* | 0,142 ± 0,027 | 0,333 ± 0,053 |
| OE *VIM1-2* | 0,132 ± 0,020 | 0,300 ± 0,046 |
| WT *vim1* | 0,122 ± 0,018 | 0,310 ± 0,068 |
| *vim1-2* | 0,132 ± 0,021 | 0,272 ± 0,039 |
| *vim1-3* | 0,137 ± 0,023 | 0,258 ± 0,040 |
| WT *NRPD2A* | 0,125 ± 0,022 | 0,296 ± 0,074 |
| *nrpd2a* | 0,115 ± 0,022 | 0,221 ± 0,053 |
| WT *RDR2* | 0,153 ± 0,045 | 0,387 ± 0,121 |
| *rdr2-5* | 0,159 ± 0,049 | 0,273 ± 0,081 |
| WT *DCL3* | 0,125 ± 0,022 | 0,296 ± 0,074 |
| *dcl3-1* | 0,164 ± 0,033 | 0,244 ± 0,041 |
| WT *AGO4* | 0,125 ± 0,022 | 0,296 ± 0,074 |
| *ago4-8* | 0,142 ± 0,034 | 0,232 ± 0,047 |
| WT *MET1* | 0,123 ± 0,028 | 0,307 ± 0,028 |
| *met1-2* | 0,110 ± 0,025 | 0,128 ± 0,038 |
| WT *CMT* | 0,131 ± 0,030 | 0,188 ± 0,044 |
| *cmt2-7* | 0,133 ± 0,028 | 0,332 ± 0,070 |
| *cmt3-11* | 0,145 ± 0,036 | 0,304 ± 0,078 |
| WT *LHP1* | 0,134 ± 0,026 | 0,276 ± 0,045 |
| *lhp1* | 0,136 ± 0,031 | 0,201 ± 0,039 |
| WT *YUC2* | 0,146 ± 0,041 | 0,337 ± 0,089 |
| *yuc2* | 0,151 ± 0,031 | 0,235 ± 0,061 |

**Supplementary Table 1: Quantification of hypocotyl length of Arabidopsis over-expression or mutant lines grown at 23°C or 29°C**

Quantification of the hypocotyl length of 4-day-old *APOLO* over-expression (OE *APOLO-1*, OE *APOLO-2*) and RNAi (RNAi *APOLO-1*, RNAi *APOLO-2*) lines, *VIM1* over-expression lines (OE *VIM1-1*, OE *VIM1-2*), *vim1* mutants (*vim1-2*, *vim1-3*), RNA-directed DNA methylation (RdDM)-related mutants (*nrpd2a, rdr2-5*, *dcl3-1*, *ago4-8*), methyltransferase mutants (*met1-2*, *cmt2-7*, *cmt3-11*), *lhp1* and *yuc2* mutants, together with their associated wild-type (WT), grown at 23°C or 29°C. Data represent average ± SD (n ≥ 60).

**Supplementary Table 2: Lists of differentially expressed genes in WT (23°C vs 29°C), *OE APOLO-1* and *vim1-3***

List of differentially expressed genes identified by RNA-seq, comparing WT plants at 23°C or 29°C, as well as *OE APOLO-1* and *vim1-3* at 23°C.

(submitted in a separate xls file)

**Supplementary Table 3: *APOLO* targets a subset of genes differentially methylated in the *vim1* mutant**

List of hypo-methylated loci in the *vim1* mutant (Stroud et al. 2013), targeted by *APOLO* (Ariel et al. 2020). The hypergeometric test was calculated considering all the Arabidopsis genes as the population size (27655 genes, Araport 11), 1974 *APOLO* targets ar the sample size, and the number of hypo-methylated genes in the respective contexts in the *vim1* mutant as the corresponding numbers of successes.

(submitted in a separate xls file)

| Primer | Sequence 5’ to 3’ | Description |
|---|---|---|
| *APOLO*.F | CTTCGAGGCGCTAAACAATC | RT-qPCR/RIP-qPCR |
| *APOLO*.R | ACAGCGGTGCCACCTATTAC | |
| *PP2A*.F | CCTGCGGTAATAACTGCATCT | RT-qPCR/RIP-qPCR |
| *PP2A*.R | CTTCACTTAGCTCCACCAAGCA | |
| *VIM1*.F | CAGTGGCATTGTCCTGATTG | RT-qPCR |
| *VIM1*.R | CACGAATCGCAGCTACAAGA | |
| *VIM1*-full.F | ATGGCGCGTGACATCCAACTCC | RT-PCR |
| *VIM1*-full.R | TCACCTGATGGTCGCAGAAAC | |
| prom*YUC2*.F | AGACAAAAGTTCAACTAGCTATACACC | ChIP-qPCR/MeDIP |
| prom*YUC2*.R | GCCAACCCGATTGATACAAA | |
| *UPAT*.F | CCCCATGTTGTTCCAAGAGT | RT-qPCR/RIP-qPCR |
| *UPAT*.R | GGTCATTGTCACCAGGTTCC | |
| exon*YUC2*.F | TTCACCTCTATTCTCGGGACA | ChIP-qPCR/ChIRP/DRIP |
| exon*YUC2*.R | CAAGAAGTGAAGTAGGGGACTGA | |
| TOPO/VIM1.F | CACCATGGCGCGTGACATC | PCR for cloning into pENTR/D-TOPO |
| TOPO/VIM1.R | TCACCTGATGGTCGCAGAAAC | |
| TOPO/PHD/RING1.F | CACCATGGCGCGTGACATC | PCR for cloning into pENTR/D-TOPO |
| TOPO/PHD/RING1.R | TCATGTAAATGCTTTATCTGG | |
| TOPO/SRA/RING2.F | CACCATGACCGAGCGTGCAAAG | PCR for cloning into pENTR/D-TOPO |
| TOPO/SRA/RING2.R | TCACCTGATGGTCGCAGAAAC | |
| TOPO/SRA.F | CACCATGACCGAGCGTGCAAAG | PCR for cloning into pENTR/D-TOPO |
| TOPO/SRA.R | TCACTCAAACAGGTCGGTGG | |
| TOPO/RING2.F | CACCATGAGAAAAGAAACTCCATC | PCR for cloning into pENTR/D-TOPO |
| TOPO/RING2.R | TCACCTGATGGTCGCAGAAAC | |
| TOPO/LHP1.F | CACCATGAAAGGGGCAAGTGGTG | PCR for cloning into pENTR/D-TOPO |
| TOPO/LHP1.R | AGGCGTTCGATTGTACTTGAGA | |
| TOPO/UPAT.F | CACCCAGCACCACACCAGG | PCR for cloning into pENTR/D-TOPO |
| TOPO/UPAT.R | GTTCAGTCACGCAGGATGG | |
| UHRF1.F | AACAGGTCTCAGGCTCAACAATGTGGATCCAGGT TCGG | PCR for cloning into pGGC000 |
| UHRF1.R | AACAGGTCTCTCTGACCGGCCATTGCCGTAGC | |
| SPACER.F | AACAGGTCTCAGGCTCAACAATGGTTAGATGTGA CAAT | PCR for cloning into pGGC000 |
| SPACER.R | AACAGGTCTCTCTGACTTAAATTCTTTCAGAAG | |
| SALK_050903_LP | AGAACTGATTCGGCAAAAAGG | Genotyping |
| SALK_050903_RP | GCGTCTCGAATCAATAATCTTG | |
| SALK_149277C_LP | GTCTGTCCTCCCAGGATTCTC | Genotyping |
| SALK_149277C_RP | GATGAGGCAACGGTTACTGAG | |

| ChIRP 3'BIO probes | | | | | |
|---|---|---|---|---|---|
| ODD probes | | EVEN probes | | EVEN LacZ probes | |
| number | sequence | number | sequence | number | sequence |
| APOLO 1 | aaccagccaatgaacagatg | APOLO 2 | gattgtttagcgcctcgaag | LacZ 1 | ccagtgaatccgtaatcatg |
| APOLO 3 | gacaagtcacacctacactc | APOLO 4 | cgagaagaactaggccaaag | LacZ 5 | aatgtgagcgagtaacaacc |
| APOLO 5 | agttccaaggaatccatacc | APOLO 6 | tgaagactaccttacataga | LacZ 7 | aataattcgcgtctggcctt |
| APOLO 7 | gacagcggtgccacctatta | APOLO 8 | acaaggaacttccaaccaaa | LacZ 9 | aattcagacggcaaacgact |
| APOLO 9 | ttacaacaagccactccgta | APOLO 10 | caacatctcgtcaaccacat | LacZ 11 | atcttccagataactgccgt |
| APOLO 11 | accaaaacaaccaaaatttt | APOLO 12 | cgaaactaaaacaaagaagc | LacZ 13 | gctgatttgtgtagtcggtt |
| APOLO 13 | ccttacaaacagagcaaagt | APOLO 14 | ggaaataacaaggcaaaaca | LacZ 15 | aactgttacccgtaggtagt |
| APOLO 15 | ggaagcaaagcccaaaggaa | APOLO 16 | ccgacgattaaaaggataat | LacZ 17 | tttcgacgttcagacgtagt |
| APOLO 17 | caagtaaccccagaaaacta | APOLO 18 | gaaatacaaagccggcggtt | LacZ 19 | accattttcaatccgcacct |
| APOLO 19 | gattccggtgaaatacaagg | APOLO 20 | tcgtctgaaagtttattata | LacZ 21 | ttcatcagcaggatatcctg |

**Supplementary Table 4: List of primers used in this study**
